# Phenotypic plasticity triggers rapid morphological convergence

**DOI:** 10.1101/2021.05.25.445642

**Authors:** José M. Gómez, Adela González-Megías, Eduardo Narbona, Luis Navarro, Francisco Perfectti, Cristina Armas

## Abstract

Phenotypic convergence, the independent evolution of similar traits, is ubiquitous in nature, happening at all levels of biological organizations and in most kinds of living beings. Uncovering its mechanisms remains a fundamental goal in biology. Evolutionary theory considers that convergence emerges through independent genetic changes selected over long periods of time. We show in this study that convergence can also arise through phenotypic plasticity. We illustrate this idea by investigating how plasticity drives *Moricandia arvensis*, a mustard species displaying within-individual polyphenism in flowers, across the morphological space of the entire Brassicaceae family. By compiling the multidimensional floral phenotype, the phylogenetic relationships, and the pollination niche of over 3000 Brassicaceae species, we demonstrated that *Moricandia arvensis* exhibits a plastic-mediated within-individual floral disparity greater than that found not only between species but also between higher taxonomical levels such as genera and tribes. As a consequence of this divergence, *M. arvensis* moves outside the morphospace region occupied by its ancestors and close relatives, crosses into a new region where it encounters a different pollination niche and converges phenotypically with distant Brassicaceae lineages. Our study suggests that, by inducing phenotypes that explore simultaneously different regions of the morphological space, plasticity triggers rapid phenotypic convergence.

## Introduction

Phenotypic convergence, the independent evolution of similar traits in different evolutionary lineages, is ubiquitous in nature, happening at all levels of biological organizations and in most kinds of living beings (*1–3*). Convergent evolution plays a fundamental role in how evolutionary lineages occupy the morphological space (*2, 4*). The expansion of lineages across the morphological space is a complex process resulting from the ecological opportunities emerging when species enter into different regions of the ecospace and face new ecological niches (*5, 6*). When this occurs, divergent selection on some phenotypes makes lineages to diversify phenotypically, boosting morphological disparity, triggering a morphological radiation and eventually filling the morphospace (*7, 8*). Because the ecological space saturate as lineages diversify (*9*), unoccupied regions become rare in highly diversified lineages (*10*). Under these circumstances, entering into a new region usually entails sharing it with other species exploiting the same ecological niche (*2, 10, 11*). In this situation, independent lineages tend to evolve similar phenotypes through convergent evolution (*2, 4*). In diversified lineages occupying a saturated morphospace, divergent and convergent evolution are ineludibly connected (*10, 12*), and both processes contribute significantly to shape the geometry of the morphospace occupation (*4, 11*).

Uncovering the mechanisms triggering convergence remains a fundamental goal in biology. Evolutionary theory shows that convergent phenotypes emerge from several genetic mechanisms, such as independent mutations or gene reuse in different populations or species, polymorphic alleles, parallel gene duplication, introgression or whole-genome duplications, that are selected over long periods of time (*13–15*). Under these circumstances, the origin of morphological convergence is mostly slow, occurring over evolutionary time and associated with multiple events of speciation and cladogenesis (*11*). It is increasingly acknowledged, however, that phenotypic plasticity might elicit the emergence of novel phenotypes with new adaptive possibilities, which may be beneficial in some contexts (*16, 17*). Under these circumstances, plasticity may behave as a facilitator for evolutionary novelty and diversity, shaping the patterns of morphospace occupation (*16, 18–21*). In this study, we provide compelling evidence showing that phenotypic plasticity also plays a prominent role in the emergence of convergent phenotypes. By inducing the production of several phenotypes, plasticity may cause the species to explore different regions of the morphospace almost simultaneously (*18, 19*). This opens the opportunity for plastic species to diverge from their lineages and converge with the species already located in other morphospace regions. We illustrate this idea by investigating how plasticity drives *Moricandia arvensis*, a species exhibiting extreme polyphenism in flowers (*18*), across the morphological space of the entire Brassicaceae family. *Moricandia arvensis* displays within-individual floral plasticity, with flower morphs varying seasonally on the same individual (*18*). By studying the multidimensional floral phenotypes, the phylogenetic relationships, and the pollination niches of over 3000 Brassicaceae species, we demonstrate that phenotypic plasticity makes the flowers of this mustard species to diverge from its ancestors and close relatives, to cross into a new region of the ecospace, and to converge morphologically with distant Brassicaceae lineages. This finding has great implications, suggesting that plasticity might not only promote the evolution of novelties and morphological divergence (*16, 17, 20, 21*) but can also provide an alternative explanation to the pervasiveness of convergence in nature.

## Results

### Plasticity-mediated floral disparity and divergence

Changes in temperature, radiation and water availability induce the production of different types of flowers by the same *M. arvensis* individuals; large, cross-shaped lilac flowers in spring but small, rounded, white flowers in summer (*18*). To quantify the magnitude of floral disparity between these two phenotypes of *M. arvensis*, we first assessed floral disparity for the entire mustard family. Brassicaceae is one of the largest angiosperm families, with almost 4000 species grouped in 351 genera and 51 tribes (*7, 22–24*). We determined the magnitude and extent of floral disparity among 3140 plant species (approx. 80% of the accepted species) belonging to 330 genera (94% of the genera) from the 51 tribes. Because we were interested in floral characters mediating the interaction with pollinators, we recorded for each studied species a total of 31 traits associated with pollination in Brassicaceae (Supplementary Data 1, Methods). We used the resulting phenotypic matrix to generate a family-wide floral morphospace. We first run a principal coordinate analysis (PCoA) to obtain a low-dimensional Euclidean representation of the multidimensional phenotypic similarity existing among the Brassicaceae species (*25*). Because the raw matrix was composed of quantitative, semi-quantitative and discrete variables, PCoA was based on Gower dissimilarities (*25*). We optimized this initial Euclidean configuration by running a non-metric multidimensional scaling (NMDS) algorithm with 5000 random starts (*25*). The resulting morphospace (Figure 1a) was significantly correlated with the initial PCoA configuration (*r* = 0.40, *P* < 0.0001, Mantel test) and was a good representation of the original relationship among the species (*R^2^* > 0.95, *Stress* = 0.2, Figure 1b). The distribution of the species across the morphospace was significantly associated with different pollination traits (Figure S1; Table S1). Species in the central region were mostly medium-sized plants bearing a moderate to high number of small, polysymmetric white flowers with short corolla tubes, exposed nectaries and visible sepals (Figure 1a, Figure S1). Species in the bottom right corner were small or prostrate, bearing minute flowers, many time apetalous and with just 2 or 4 stamens, whereas species located in the bottom left corner were medium-sized plants with asymmetric flowers arranged in corymbous inflorescences. Plants with yellow flowers were located in the right region of the morphospace. In contrast, large plants with strongly tetradynamous androceum and large, veined, dissymmetrical to asymmetrical, pink to blue flowers with concealed nectaries, long corolla tubes and bullseyes were located in the upper left region (Figure 1a, Figure S1). *Moricandia arvensis*, when blooming in spring (Figure 1c), occupies this later peripheral region of the morphospace, close to other *Moricandia* species (purple dots in Figure 1a). However, during summertime, the individuals of *M. arvensis* are shorter and produce fewer, much smaller flowers with white, unveined and rounded corollas with overlapped petals and green sepals that are mostly arranged alone the floral stems (Figure 1d) (*18*). Due to this radical phenotypic change, the summer phenotype of *M. arvensis* was located in a different, more central position of the floral morphospace (Figure 1a), far away from the region occupied by the *Moricandia* species. As a consequence of this jump, the morphological disparity between the spring and summer phenotypes of *M. arvensis*, calculated as their distance in the morphospace (*26*), was very high (0.264). In fact, it was much higher than the average pairwise disparities among all studied Brassicaceae species (0.155 ± 0.090, mean ± s.e.m., 4,912,545 pairwise disparities) and almost 50% of the largest observed disparity (0.55) (Table S4). This outcome suggests that phenotypic plasticity prompts *M. arvensis* to explore two distant regions of the Brassicaceae floral morphospace simultaneously.

**Figure 1.**
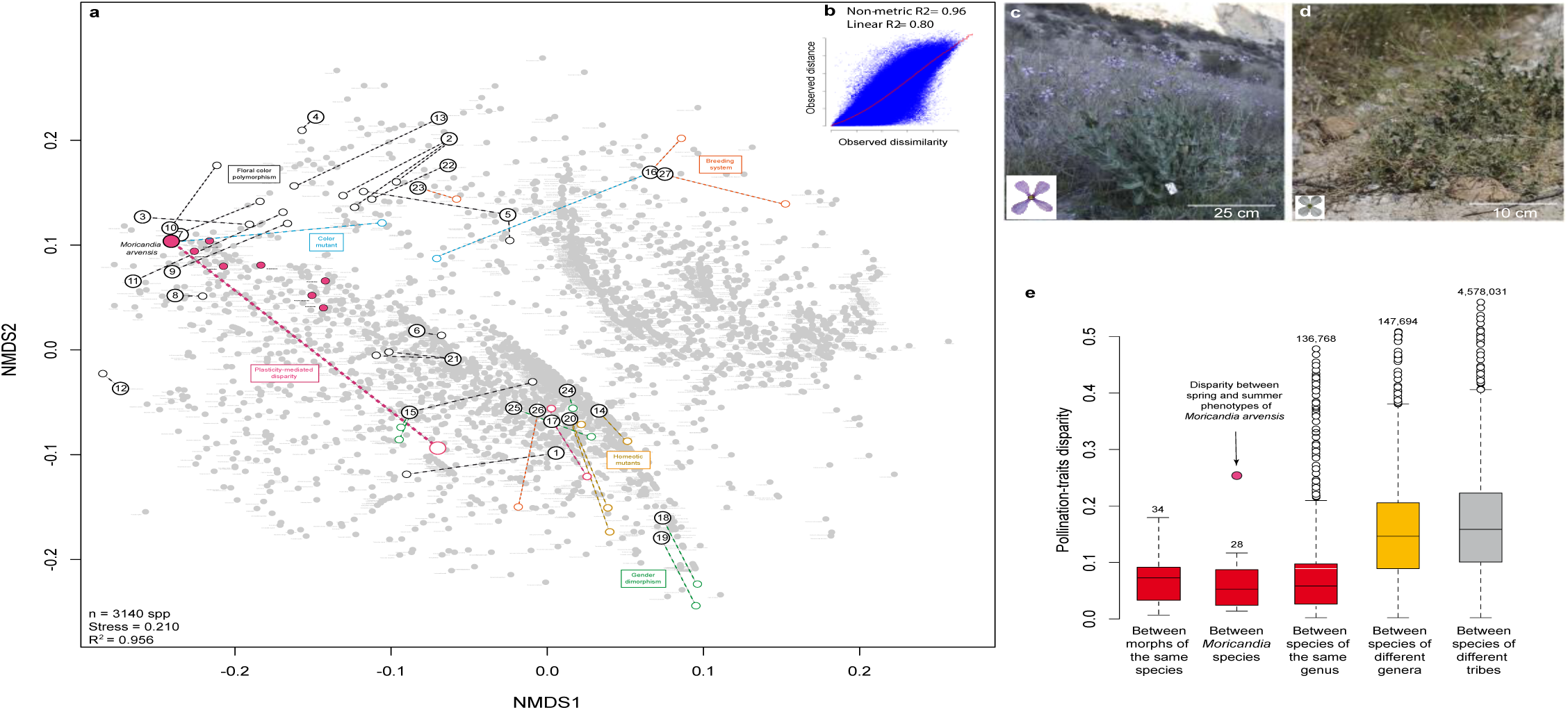
Plasticity-mediated floral disparity. (**A**) Floral morphospace of the Brassicaceae, showed as the projection of 31 traits recorded in 3140 species onto two NMDS axes. The position of the spring and summer phenotypes of *Moricandia arvensis* is linked by a thick lilac dashed line. We have also indicated the movements across this morphospace of several species changing their phenotypes due to floral colour polymorphism (black lines), single mutations in floral colour (blue lines), changes in breeding systems (orange lines), changes in gender expression (green lines), homeotic mutations (brown lines), and plasticity (lilac lines). Numbers matching species are as follow: 1-*Lobularia maritima*; 2-*Raphanus raphanistrum*; 3-*Matthiola incana*; 4-*Mathiola fruticulosa*; 5-*Erysimum cheiri*; 6-*Cakile maritima*; 7-*Matthiola lunata*; 8-*Marcus-kochia littorea*; 9-*Hesperis matronalis*; 10-*Hesperis laciniata*; 11-*Parrya nudicalis*; 12-*Streptanthus glandulosus*; 13-*Eruca vesicaria*; 14-*Capsella bursa-pastoris*; 15-*Hormathophylla spinosa*; 16-*Brassica napus*; 17-*Cardamine hirsuta*; 18-*Lepidium sisymbrioides*; 19-*Lepidium solandri*; 20-*Arabidopsis thaliana*; 21-*Boechera stricta*; 22-*Leavenworthia stylosa*; 23-*Leavenworthia crassa*; 24-*Pachycladon stellatum*; 25-*Pachycladon wallii*; 26-*Cardamine kokairensis*; 27-*Brassica rapa*. (**B**) Shepard plot showing the goodness of fit of the NMDS ordination. (**C**) *Moricandia arvensis* in spring. (**D**) *Moricandia arvensis* in summer. (**E**) Magnitude of floral disparity between different taxonomic levels of Brassicaceae species. The number above each boxplot shows the number of disparities per level. We have compared this value with the disparity between spring and summer phenotypes of *M. arvensis* (this comparison with boxplots in red is statistically significant at *P* < 0.05, in orange is marginally significant at *P* < 0.1, and in grey is non-significant).

To know how intense is the plasticity-mediated *M. arvensis* disparity, we compared its value with the disparity values observed at different taxonomic levels within Brassicaceae. At the lowest level, discrete changes in pollination traits have been reported between individuals of the same species. In some species, this intraspecific phenotypic change is stable, like the gender polymorphism (*27, 28*) or the adaptive floral colour polymorphism exhibited as a response to the selective pressures exerted by certain pollinators (*29, 30*). In other species, discrete phenotypic changes, although affecting pollination traits, seem to be just the consequence of some singular and often unstable mutations affecting floral colour (*31*), the production of cleistogamous flowers (*32*) or changes in the expression of homeotic genes that modify the formation of the floral organs (*33, 34*). We compiled information on the phenotypes of the different morphs in 34 polymorphic species and calculated their values of intraspecific disparities (Figure 1a, Supplementary Data 2). Although several polymorphic species showed considerable values of between-morph disparity, they were significantly smaller than the disparity between spring and summer floral phenotypes of *M. arvensis* (*Z-*score = 5.06, *P* < 0.0001, Figure 1e, Table S2). We subsequently tested at what taxonomic level of Brassicaceae the disparity was equivalent to the plasticity-mediated disparity observed in *M. arvensis*. For this, we calculated the floral disparity between pair of species belonging to the genus *Moricandia*, the same genus, the same tribe, and different tribes (Methods). The plasticity-mediated disparity of *M. arvensis* was significantly higher than the disparity existing between the *Moricandia* species (0.057 ± 0.033, mean ± 1 s.e.m., *Z*-score = 6.27, *P* < 0.0001) and between the species belonging to the same genus (0.069 ± 0.055, *Z*-score = 3.51, *P* < 0.0002). It was marginally different from the disparity existing between species of different genera but the same tribes (0.150 ± 0.085, *Z*-score = 1.34, *P* = 0.089) and it was statistically similar to the disparity occurring between species belonging to different tribes (0.167 ± 0.087, *Z*-score = 1.11, *P* = 0.133, Figure 1e). These findings suggest that phenotypic plasticity allows *M. arvensis* individuals to jump in the morphospace longer distances than those granted by some macroevolutionary processes.

We explored whether plasticity-mediated disparity may cause evolutionary divergence by calculating the disparity of *M. arvensis* spring and summer phenotypes to their phylogenetic ancestors. We retrieved 80 partial phylogenies from the literature and online repositories (Methods), and assembled them into a supertree comprising 1876 taxa with information on their floral phenotype. We then projected this supertree onto the morphospace to get a family-wide phylomorphospace. We did not find evidence of phylogenetic constraints on morphospace occupation since there was not significant phylogenetic signal for the position occupied by each species (Multivariate *Mantel* test=0.005, *P* = 0.34). The family-wide phylomorphospace was very tangled (Figure 2a), with 492,751 intersections among lineages, suggesting the presence of many events of floral divergence and convergence in the evolution of Brassicaceae pollination traits (*11*). To calculate the disparity of the *M. arvensis* floral phenotypes to their ancestor, because these analyses are sensitive to the tree topology and the inferred branch lengths (*26*), we used four independent, time-calibrated phylogenies that included this species (Methods). The results were consistent across phylogenies (Figure 2b,c; Tables S3). The spring phenotype did not significantly diverge neither from the most recent common ancestor (MRCA) of *Moricandia* (*Z*-score = 0.36, *P* = 0.36) nor from its direct ancestor (*Z*-score = −1.24, *P* = 0.108). In contrast, the summer phenotype of *M. arvensis* diverged significantly both from *Moricandia* MRCA (*Z*-score = 2.48, *P* = 0.007) and from its direct ancestor (*Z*-score = 1.77, *P* = 0.038). Hence, the summer phenotype explores a region of the floral morphospace located out of its phylogenetic clade range (Figure 2b). The ancestral disparity of the summer phenotype was even significantly higher than the ancestral disparity of most other Brassicaceae species (Figure 2c). These findings suggest that phenotypic plasticity causes the appearance of a novel phenotype that diverges radically from its ancestors.

**Figure 2.**
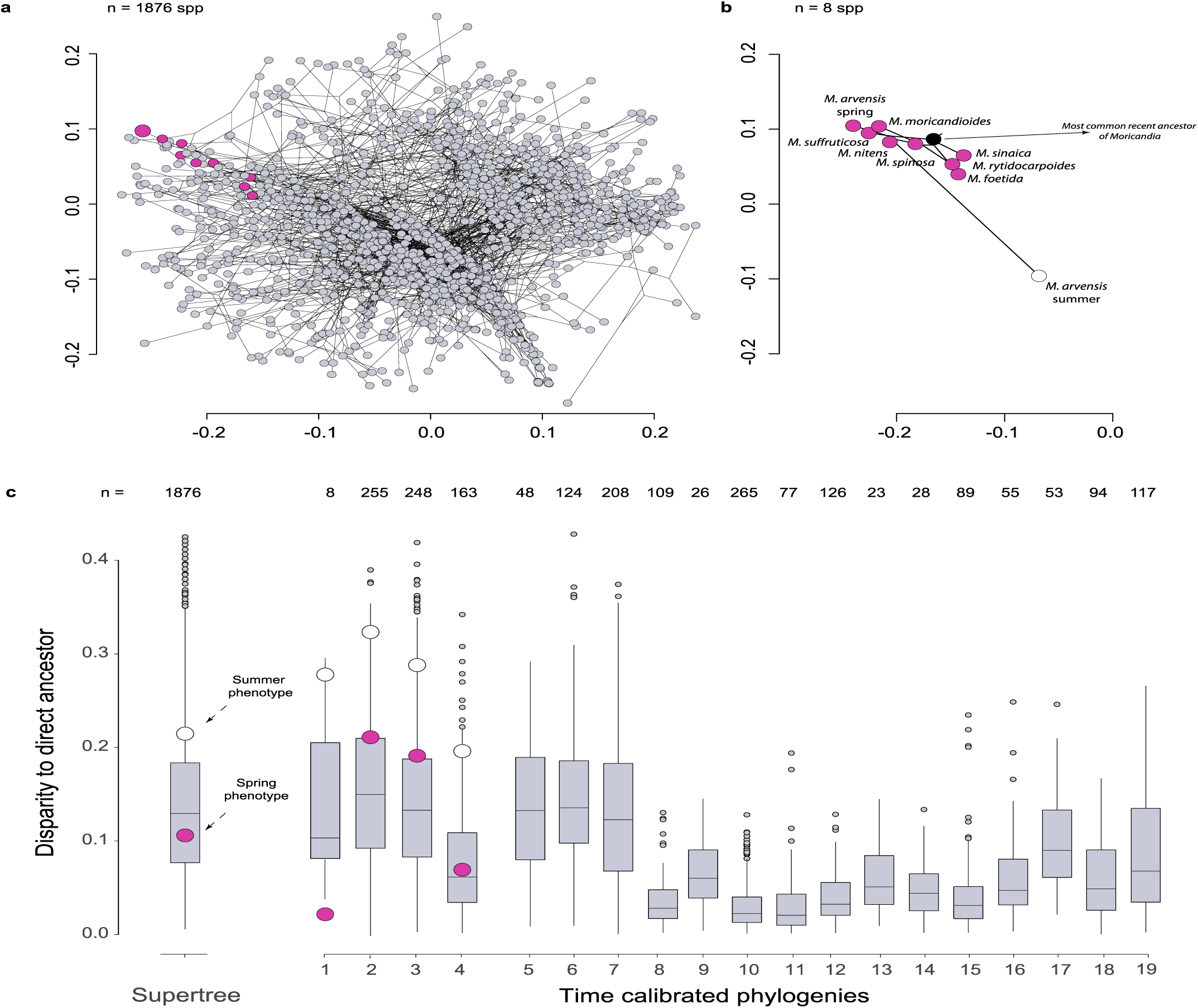
Phylogenetic-mediated floral divergence. (**A**) Floral phylomorphospace using the supertree that includes 1876 Brassicaceae species. (**B**) Phylomorphospace considering only the eight *Moricandia* species, using the Perfectti et al.’s phylogeny (phylogeny # 1 in Table S8). (**C**) Floral disparity to the nearest ancestor, according to the supertree and 18 time-calibrated phylogenies (phylogeny codes in Table S8). We show the disparity between the two *M. arvensis* phenotypes and their direct ancestor (spring: lilac dots; summer: white dots) in those phylogenies that include *Moricandia*. We also show the disparities to their direct ancestors of those Brassicaceae species included in time-calibrated phylogenies of more than 45 species.

### Plastic shifts in pollination niches

Evolutionary divergence is mostly associated with the occupation of new ecological niches (*2, 5*). Shifts between pollination niches are an important factor driving diversification in angiosperms (*35*), including Brassicaceae (*36, 37*). We investigated whether the plasticity-mediated jump of *M. arvensis* across the floral morphospace implicated the exploration of new pollination niches. We compiled a comprehensive database comprising 456,031 visits done by over 800 animal species from 19 taxonomical orders, 276 families and 43 functional groups to 554 Brassicaceae species of 39 tribes (Methods, Supplementary Data 3). Afterwards, we identified the pollination niches of these Brassicaceae plants and determined the niche of each *M. arvensis* floral phenotype by means of bipartite modularity, a complex network tool that identifies the set of plants interacting with similar groups of pollinators (*18*). This analysis showed that the network was significantly modular (*Modularity* = 0.385, *P* < 0.0001) and identified eight different pollination niches associated with different groups of pollinators (Figure 3a) located in different regions of the morphospace (Figure 3b, *F* = 44.4, *P* < 0.001, *R^2^* = 0.39, Adonis test; Table S4).

**Figure 3.**
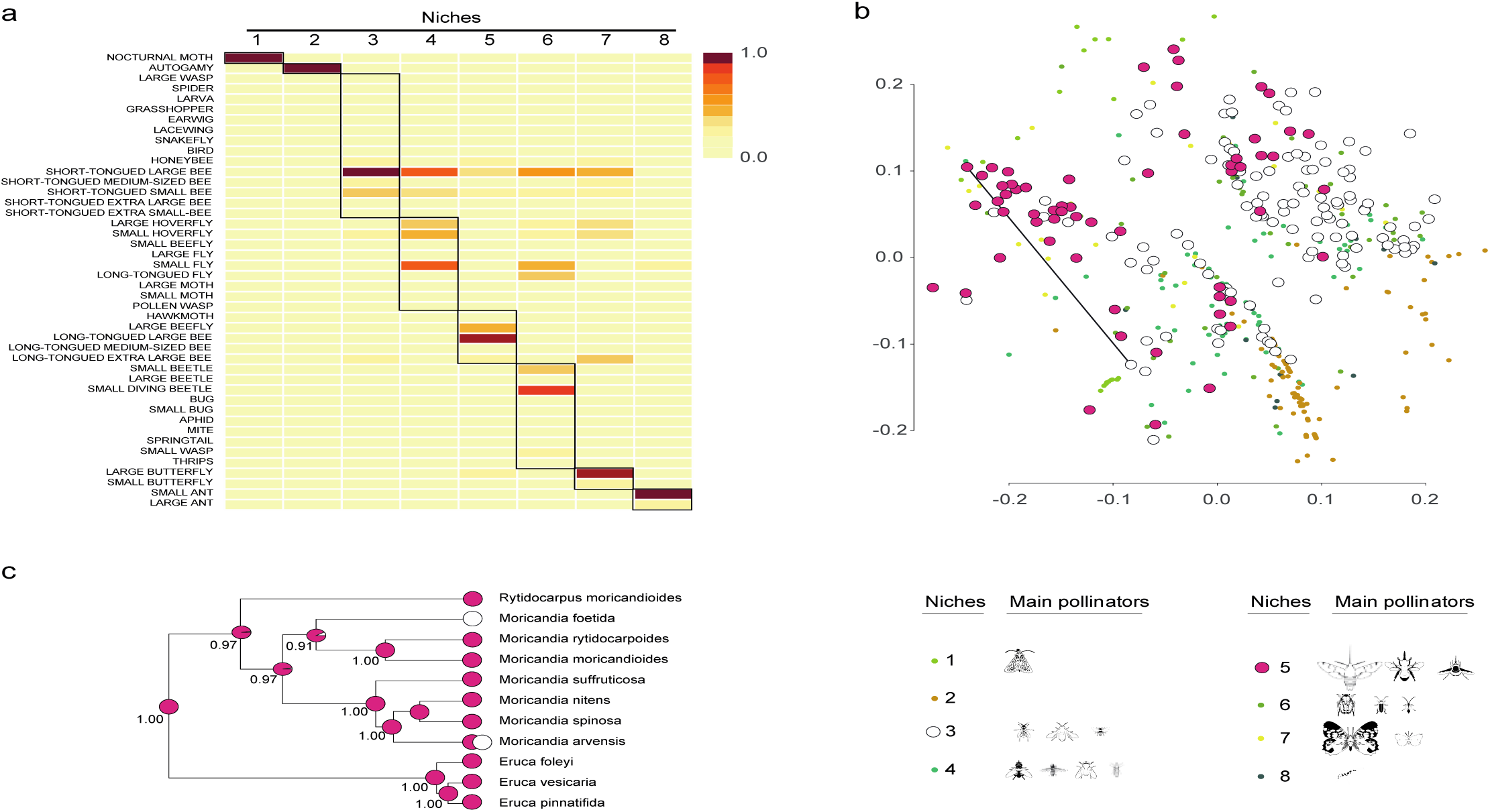
Plasticity-mediated changes in pollination niches. (**A**) Outcome of the modularity analysis showing the number of pollination niches inferred, the among-niche differences in relative frequency of each pollinator functional group, and the pollinator functional groups defining the niches (n = 511 Brassicaceae species). (**B**) Morphospatial distribution of the eight pollination niches detected in Brassicaceae. Insect silhouettes were drawn by Divulgare (www.divulgare.net) under a Creative Commons license (http://creativecommons.org/licenses/by-nc-sa/3.0). (**C**) Estimate of the ancestral pollination niche of the *Moricandia* lineage using a stochastic character mapping inference analysis. The numbers underneath each ancestral node indicate the posterior Bayesian probability of belonging to pollination niche 5.

Because different insects visited *M. arvensis* in spring and summer (Table S5), this plant species shifted between pollination niches seasonally (Figure 3b). During spring, *M. arvensis* belonged to a niche where most frequent pollinators were long-tongued bees, beeflies, and hawkmoths (pollination niche 5 in Figure 3a) (*18*). This pollination niche was also shared by the other *Moricandia* species (Figure 3c). In contrast, during summer *M. arvensis* belonged to a niche dominated by short-tongued bees (pollination niche 3 in Figure 3a). This niche shift was substantial. In fact, the overlap between the spring and summer pollinator niches of *M. arvensis* (*Czekanowski* overlap index = 0.35) was significantly lower than the overlap between congeneric species of Brassicaceae (0.57 ± 0.42, *Z*-score = −0.51, *P* = 0.003). This shift even entailed the divergence from the ancestral niche of the *Moricandia* lineage (pollination niche 5 according to a stochastic character mapping inference, Figure 3c). The within-individual floral plasticity allows *M. arvensis* to exploit a pollination niche that differs markedly from that exploited by its closest relatives and that have largely diverged from the ancestral niche.

### Plasticity-mediated floral convergence

A common consequence of adaptation to the same niche is convergent evolution (*1, 2, 4*). We explored the possibility of convergent evolution of *M. arvensis* with other Brassicaceae sharing either the spring niche (pollination niche 5) or the summer niche (pollination niche 3). We first checked for the occurrence of convergence among species belonging to these pollination niches. Because these analyses are extremely sensitive to the inferred branch lengths, we explored morphological convergence using three time-calibrated large (> 150 spp) phylogenies that included *M. arvensis* (Methods). We tested for the occurrence of floral convergence between the species belonging to each of those two pollination niches using three methods: the angle formed by the phenotypic vectors connecting the position in the floral morphospace of each pair of species with that of their most recent common ancestor (*38*), the difference in phenotypic distances between convergent species and the maximum distances between all other lineages (*39*), and the phenotypic similarity of the allegedly convergent species penalized by their phylogenetic distance (*Wheatsheaf* index) (*40*). The three methods gave similar results, indicating that floral convergence was frequent among the species belonging to any of the two studied niches, irrespective of the method and the time-calibrated tree used (Table S6). These results show that, despite the rampant generalization observed in the pollination system of Brassicaceae, species interacting with similar pollinators converge phenotypically.

Once we determined the occurrence of convergence in these two pollination niches, we assessed whether plasticity caused the evolution of morphological convergence in *M. arvensis*. To do so, we first assessed the convergence region of *Moricandia*, the region that includes the lineages converging morphologically to the *Moricandia* lineage. We found that this region included most species of *Moricandia*, the spring phenotype of *M. arvensis*, and several clades belonging to disparate tribes that interact with pollination niche 5, but excluded the summer phenotype of *M. arvensis* (Figure 4, Table S7). Afterwards, we checked whether any of the two *M. arvensis* floral phenotypes entered the region of the phylomorphospace defined by their pollination niches. We used the C5 index, defined as the number of lineages that cross into the morphospace region of interest from outside^39^. This index detected between two and six convergent events towards pollination niche 5 depending on the phylogeny used (blue arrows in Figure 4a-c), but none was associated with the spring phenotype of *M. arvensis*. In contrast, the C5 index consistently detected that the summer phenotype of *M. arvensis* has converged with the species belonging to the pollination niche 3 (red arrow in Figure 4d-f). Altogether, these analyses suggest that, whereas the spring phenotype did not show any evidence of convergence, the summer phenotype of *M. arvensis* has converged with other distant Brassicaceae exploiting the same pollination niche.

**Figure 4.**
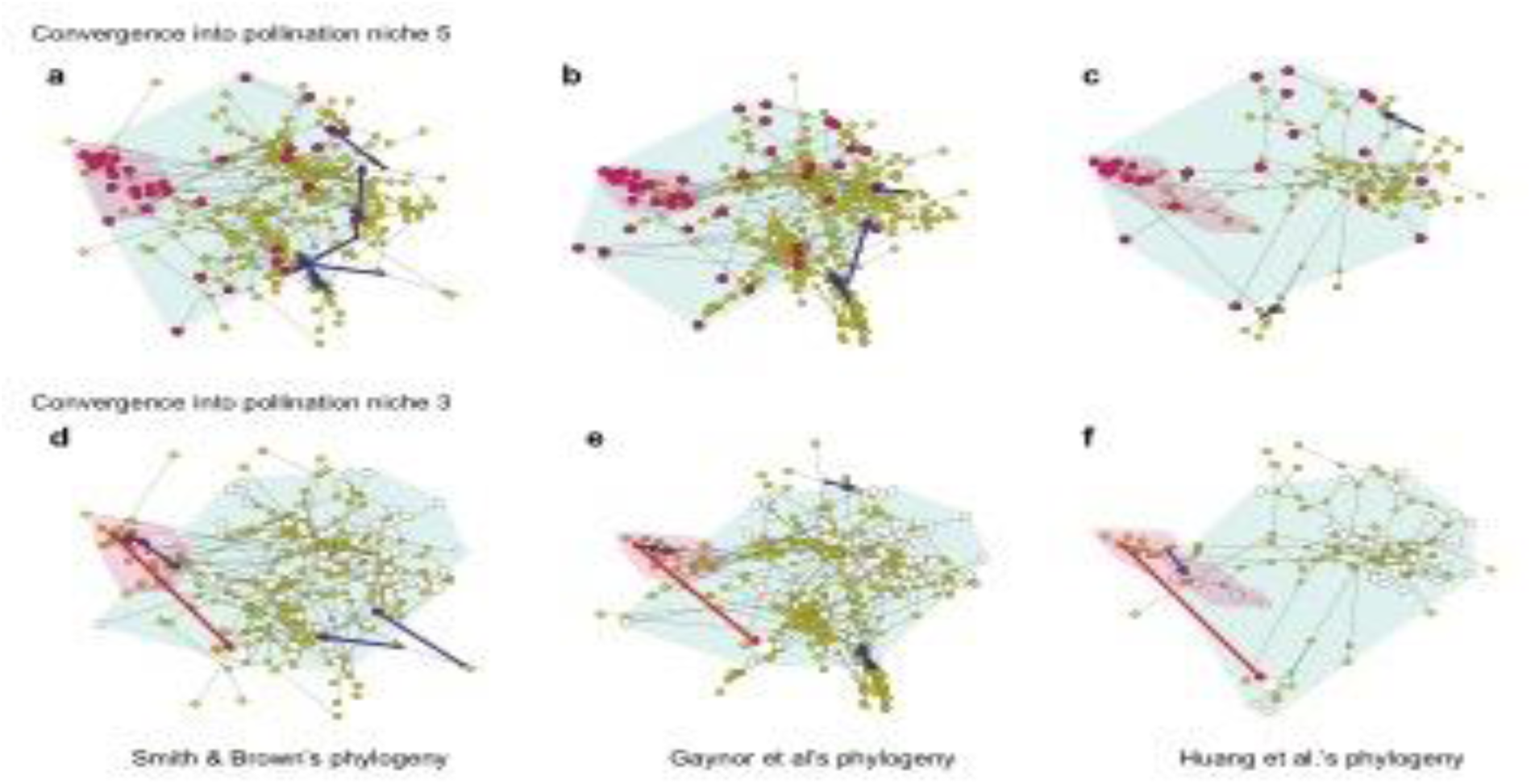
Plasticity-mediated floral convergence. Convergent lineages crossing into the region of the morphospace delimited by the pollination niche of the *M. arvensis* during spring (the shade convex hull) according to (**A**) Smith & Brown’s phylogeny, (**B**) Gaynor et al.’s phylogeny, and (**C**) Huang et al.’s phylogeny (phylogenies 2-4, respectively, in Table S8). Convergent lineages crossing into the region of the morphospace delimited by the pollination niche of the *M. arvensis* during summer (the shade convex hull) according to (**D**) Smith & Brown’s phylogeny, (**E**) Gaynor et al.’s phylogeny, and (**F**) Huang et al.’s phylogeny. Red arrows indicate the plasticity-mediated convergence, blue arrows the convergence events of the other lineages. The small purple area in all panels is the region of the floral morphospace that includes the lineages that have converged with the entire *Moricandia* clade according to each time-calibrated phylogeny.

## Conclusions

Convergent selection exerted by efficient pollinators causes the evolution of similar suites of floral traits in different plant species (*41–44*). Our study shows that plasticity can promote the rapid convergent evolution of floral traits, providing an additional explanation about how pollination syndromes may evolve. Under this idea, changes in floral traits precede shifts in pollinators, as frequently observed in generalist systems (*37, 45*). This may explain why many pollination systems are evolutionarily labile, undergoing frequent shifts and evolve multiple times within the same lineages by diverse evolutionary pathways (*35, 46*).

Morphological convergence is universally acknowledged to be the result of several genetic mechanisms, such as independent mutations in different populations or species, polymorphic genes or introgression (*13*). We provide in this study compelling evidence suggesting that morphological convergence may also arise as a consequence of phenotypic plasticity. The role of plasticity as a mechanism favouring quick responses of organisms to novel and rapidly changing environments is already beyond doubt (*17, 21, 47, 48*). Its evolutionary consequences are more debated though (*20, 21, 49, 50*). The ‘plasticity-led evolution’ hypothesis states that selection acting on a plastic lineage may either boost its environmental sensitivity and trigger the origin of polyphenisms or alternatively may promote the loss of plasticity and the canalization of the new phenotype through genetic assimilation (*21, 49*). The related ‘flexible stem’ hypothesis of adaptive radiation suggests that when a plastic lineage repeatedly colonizes similar niches, the multiple phenotypes fixed by genetic assimilation could converge among them giving rise to a collection of phylogenetically related convergent morphs (*16, 50, 51*). Our comprehensive study complements these hypotheses by suggesting that plasticity-mediated convergence may even evolve without the existence of basal flexible lineages. Rather, it can occur when plasticity evolving in otherwise non-plastic lineages promotes the colonization of a niche previously occupied by unrelated species. Under these circumstances, contrary to what it is predicted by the previous hypotheses, plasticity-mediated convergence is not circumscribed to phylogenetic-related species arising from a common stem lineage. This overlooked role of phenotypic plasticity may contribute to explain the ubiquity of morphological convergence in nature.

## Materials and Methods

### Floral traits

We recorded from the literature 31 floral traits in 3140 Brassicaceae plant species belonging to 330 genera and 51 tribes (Supplementary Data 1). All these traits have been proven to be important for the interaction with pollinators (Table S8). These traits were: (1) Plant height; (2) Flower display size; (3) Inflorescence architecture; (4) Presence of apetalous flowers; (5) Number of symmetry axes of the corolla; (6) Orientation of dominant symmetry axis of the corolla; (7) Corolla with overlapped petals; (8) Corolla with multilobed petals; (9) Corolla with visible sepals; (10) Petal length; (11) Sepal length; (12) Asymmetric petals; (13) Petal limb length; (14) Length of long stamens; (15) Length of short stamens; (16) Stamen dimorphism; (17) Tetradynamous condition; (18) Visible anthers; (19) Exserted stamens; (20) Number of stamens; (21) Concealed nectaries; (22) Petal carotenoids; (23) Petal anthocyanins; (24) Presence of bullseyes; (25) Presence of veins in the petals; (26) Coloured sepals; (27) Relative attractiveness of petals versus sepals; (28) Petal hue; (29) Petal colour as b CIELAB; (30) Sepal hue; (31) Sepal colour as b CIELAB. A detailed definition and description of these traits and their states is provided in Key Resource Table 1, whereas the original references used to determine the states of each trait per plant species is provided in Supplementary Data 1.

### Family-wide floral morphospace

Using the original multidimensional trait-species matrix, we built a floral morphospace. For this, we reduced the high-dimensional matrix of floral traits to a two-dimensional space using an ordination technique (*25*). Because the set of floral traits included in this study were quantitative, semi-quantitative and qualitative, we used ordination techniques based on dissimilarity values. For this, we first constructed a pairwise square distance matrix of length equal to the number of Brassicaceae species included in the analysis (n = 3140). We used the Gower distance, the number of mismatched traits over the number of shared traits. This dissimilarity index is preferable to the raw Euclidean distance when there are discrete and continuous traits co-occurring in the same dataset (*52*).

We reduced the dimensionality of this phenotypic matrix by projecting it in a two-dimensional space. For this, to ensure an accurate description of the distribution of the species in the morphospace, we first run a principal coordinate analysis (PCoA), a technique providing a Euclidean representation of a set of objects whose relationship is measured by any dissimilarity index. We corrected for negative eigenvalues using the Cailliez procedure (*25*). Afterwards, we used this metric configuration as the initial configuration to run a non-metric multidimensional scaling (NMDS) algorithm (*25*), a method that will further optimise the sample distribution so as more variation in species composition is represented by fewer ordination axes. Unlike methods that attempt to maximise the variance or correspondence between objects in an ordination, NMDS attempts to represent, as closely as possible, the pairwise dissimilarity between objects in a low-dimensional space. NMDS is a rank-based approach, where the original distance data is substituted with ranks, preserving the ordering relationships among species (*25*). Objects that are ordinated closer to one another are likely to be more similar than those further apart (*53*). This method is more robust than distance-based methods when the original matrix includes variables of contrasting nature. However, NMDS is an iterative algorithm that can fail to find the optimal solution. We decreased the potential effect of falling in local optima by running the analysis with 5000 random starts and iterating each run 1 x 10^6^ times (*54*). The NMDS was run using a monotone regression minimizing the Kruskal’s stress-1 (*55, 56*), and compared each solution using Procrustes analysis, retaining that with the lowest residual. Because many species did not share trait states, a condition complicating ordination, we used *stepacross* dissimilarities, a function that replaces dissimilarities with shortest paths stepping across intermediate sites while regarding dissimilarities above a threshold as missing data (*57*). Furthermore, we used weak tie treatment, allowing equal observed dissimilarities to have different fitted values. The scores of the species in the final ordination configuration were obtained using weighted averaging. We checked if the reduction in dimensionality maintained the between-species relationship by checking the stress of the resulting ordination and finding goodness of fit measure for points in nonmetric multidimensional scaling (*54*). Both PCoA and NMDS ordinations were done using the R package *vegan* (*58*) and *ecodist* (*59*). It is important to note that, although the transfer function from observed dissimilarities to ordination distances is non-metric, the resulting NMDS configuration is Euclidean and rotation-invariant (*60*).

### Morphological Disparity

Because we were interested in describing the position of the species in the floral morphospace, we calculated the morphological disparity using indices related to the distance between elements (*26, 61*). We first determined the absolute position of each of the Brassicaceae species in the morphospace by calculated their Euclidean distance with the overall centroid of the morphospace (*61*). The disparity between the spring and summer phenotype of *M. arvensis* was also calculated as their Euclidean distance in the floral morphospace. We then calculated the pairwise disparities between all species included in our analysis, between the different morphs of the polymorphic species considered here (Supplementary Data 2), between the species of the genus *Moricandia*, between species of the same genus, between species of different genera but same tribe and between species of different tribes. These disparity values were calculated using the function *dispRity* of the R package *dispRity* using the command *centroid* (*62*). We checked whether the disparity between spring and summer *M. arvensis* phenotypes was significantly different from the disparities of each of these sets of species using *Z*-score tests.

### Family-wide phylogeny

We retrieved 80 phylogenetic trees from the literature and from the online repositories TreeBase (Table S9). All trees were downloaded in nexus format. The taxonomy of the species included in each tree was checked and updated using the species checklist with accepted names provided by Brassibase (https://brassibase.cos.uni-heidelberg.de/) (*7, 23, 63*). All trees were converted to TreeMan format (*64*) and concatenated into a single TreeMen file that was then converted into a multiPhylo class. Afterward, we estimated a supertree from this set of trees. Because trees did not share the same taxa, we used the Matrix representation parsimony method (*65*). To make this supertree more accurate, it was re-constructed using as backbone phylogeny the tree provided by Walden et al. (*7*). We removed from the supertree those species without information on floral phenotype, resulting in a tree with 1876 taxa. Because the original trees used to assemble this supertree where very heterogeneous, this supertree was not dated. We finally rooted the supertree using several species belonging to the sister families Capparaceae and Cleomaceae (*66*). All phylogenetic manipulations were performed using the R libraries *treebase* (*67*), *ape* (*68*), *treeman* (*64*), *phangorn* (*69*) and *phytools* (*70*).

We tested whether the position of the Brassicaceae species in the morphospace was associated with the phylogenetic relationship by assessing the phylogenetic signal of the morphospace position. This analysis was performed by means of a multivariate Mantel test, using the pairwise disparity (the Euclidean distance between species in the family-wide morphospace) as a morphological distance and the patristic distances between pairs of tips of the supertree as the phylogenetic distance (*71*). The correlation method used was Pearson and the statistical significance was found after bootstrapping 999 times the analysis (*25*). The test was done using the R libraries *vegan* (*58*) and *ecodist* (*59*).

### Family-wide phylomorphospace

We reconstructed a family-wide phylomorphospace by projecting the phylogenetic relationships provided by the supertree into the floral morphospace. The ancestral character estimation of morphospace coordinate values for each internal tree node was done using maximum likelihood. For this, we used the function *fastAnc* in *phytools*. This function performs fast estimations of the ML ancestral states for continuous traits by re-rooting the tree at all internal nodes and computing the contrasts state at the root each time (*70*).

We counted the number of intersections between lineages as a measurement of the disorder of the phylomorphospace and evidence of the mode of evolution of the phenotypes (*11*). For this, we used R codes provided in Ref *11*. We compared the observed number of crossings with those expected under several modes of evolution. For this, we counted the number of intersections in 10 simulated sets of species with floral phenotypes following Brownian Motion, Ornstein Uhlenbeck and Early Burst modes of evolution. All simulations were done using as backbone tree the family-wide supertree and considering 1875 species, and by means of the command mvSIM in *mvMORPH* (*72*).

### Morphological divergence of the plastic phenotypes

Divergence in floral phenotype was estimated by calculating the disparity of *Moricandia arvensis* and the rest of Brassicaceae species from their ancestors. We first determined the floral phenotype of the Most Recent Common Ancestor (MRCA) using the projection of a recent time-calibrated phylogeny made for the genus *Moricandia* (*73*) into the above-described phylomorphospace. We used this phylogeny because it is the only one including all the species of the genus. Once we inferred the coordinates of the MRCA in the morphospace, we calculated the disparity of all the *Moricandia* species and the two plastic phenotypes of *M. arvensis* to it. Afterwards, we calculated the divergence of the two plastic phenotypes from the direct ancestor of *M. arvensis*. This analysis was done for the family-wide supertree and for any of the four time-calibrated phylogenies included in our dataset that had *Moricandia* species (*73–76*). In addition, we calculated the divergence from the direct ancestors of the rest of Brassicaceae species included in these four phylogenies and in the rest of the time-calibrated trees included in our dataset (Table S9). All floral divergences were calculated using the command *ancestral.dist* of the function *dispRity* in the R package *dispRity* (*62*).

### Pollinator Database

We have compiled a massive database including 21,212 records comprising 455,014 visits done by over 800 animal species from 19 taxonomical orders, 276 families and 43 functional groups to 554 Brassicaceae species belonging to 39 tribes (Supplementary Data 3). Information is coming from literature, personal observation, online repositories and personal communication of several colleagues. The source of information is indicated in the database (Supplementary Data 3, Table S10). In those species studied by us (coded as UNIGEN data origin in the Supplementary Data 3), we conducted flower visitor counts in 1-16 populations per plant species. We visited the populations during the blooming peak, always at the same phenological stage and between 11:00 am and 5:00 pm. In these visits, we recorded the insects visiting the flowers for two hours without differentiating between individual plants. Insects were identified in the field, and some specimens were captured for further identification in the laboratory. We only recorded those insects contacting anthers or stigma and doing legitimate visits at least during part of their foraging at flowers. We did not record those insects only eating petals or thieving nectar without doing any legitimate visit. The information obtained from the literature and online repositories (coded as LITERATURE data origin in the Supplementary Data 3) includes records done during ecological studies, taxonomical studies and naturalistic studies. The reference of every record is included in the dataset. The plant species included in our network do not coexist, implying that this is a clade-oriented network rather than an ecological network (*77*).

### Spatial distribution of pollinator groups

We tested the autocorrelation across the morphospace in the abundance of the functional groups using a multivariate Mantel test. The correlation method used was Pearson, and the statistical significance was found after bootstrapping 999 times the analysis (*25*). The test was done using the R libraries *vegan* (*58*).

### Pollination niches

In plant species interacting with a diverse assemblage of pollinators, like those included in this study, many pollinator species interact with the flowers in a similar manner, have similar effectiveness and exert similar selective pressures and are thus indistinguishable for the plant (*46, 78*). These pollinators are thus grouped into functional groups, which are the relevant interaction units in generalised systems (*46, 78, 79*). We thereby grouped all pollinators visiting the Brassicaceae species using criteria of similarity in body length, proboscis length, morphological match with the flower, foraging behaviour, and feeding habits (*46, 78, 79*). Table S11 describes the 43 functional groups used in this study. Supplementary Data 4 shows the species with an autogamous pollination system.

We determined the occurrence of different pollination niches in our studied populations and seasons using bipartite modularity, a complex-network metric. Modularity has proven to be a good proxy of interaction niches both in ecological networks, those included coexisting species or population, as well as in clade-oriented network, those including species with information coming from disparate and contrasting sources (*77*). We constructed a weighted bipartite network, including pollinator data of four populations during the spring and summer flowering. In this network, we pooled the data from the different individuals in a population and did not consider the time difference involved in sampling across different species. We removed all plant species with less than 20 visits. We subsequently determined the modularity level in this weighted bipartite network by using the QuanBiMo algorithm (*80*). This method uses a Simulated Annealing Monte-Carlo approach to find the best division of populations into modules. A maximum of 10^10^ MCMC steps with a tolerance level = 10^-10^ was used in 100 iterations, retaining the iterations with the highest likelihood value as the optimal modular configuration. We tested whether our network was significantly more modular than random networks by running the same algorithm in 100 random networks, with the same linkage density as the empirical one (*81*). Modularity significance was tested for each iteration by comparing the empirical versus the random modularity indices using a *Z*-score test (*80*). After testing the modularity of our network, we determined the number of modules (*82*). We subsequently identified the pollinator functional groups defining each module and the plant species ascribed to each module. Modularity analysis was performed using the R package *bipartite* 2.0 (*83*). We quantified the niche overlap between all pair of Brassicaceae species using the Czekanowski index of resource utilization, an index that measures the area of intersection of the resource utilization histograms of each species pair (*84*). This index was calculated using the function *niche.overlap* in the R package *spaa* (*85*).

### Estimation of ancestral values of pollination niches

The ancestral states of the pollination niche was inferred for the *Moricandia* lineage by simulate stochastic character mapping of discrete traits with Bayesian posterior probability distribution (*86, 87*). Three models of character evolution (“ER” - Equal Rates; “SYM” – symmetric; and “ARD” - All Rates Different) were first evaluated using the *fitDiscrete* function of the R package *Geiger* (*88*). The best model was selected using the Akaike Information Criterion (AIC) and used for stochastic character mapping. The posterior distribution of the transition rate matrix was determined using a Markov chain Monte Carlo (MCMC) simulation, and the stochastic mapping was simulated 100 times. Stochastic character mapping was performed using the *make.simmap* function and a plot of posterior probabilities were mapped using the *describe.simmap* function in R package ‘*phytools* (*70*).

### Morphological convergence

To explore morphological convergence, we reconstructed the ancestral states of the species belonging to these two pollination niches and tested for each niche whether the species were morphologically more similar to each other than expected by their phylogenetic relationship (*39, 40*). We used three different approaches to detect morphological convergence, one based on comparing phenotypic and phylogenetic distances (*39*) and the other based on comparing the angles formed by two tested clades from their most recent common ancestor with the expected angle according to null evolutionary models (*38*). Because all these analyses are sensitive to the number of tips in the phylogeny and the inferred branch lengths, we tested for the occurrence of morphological convergence using three independent, time-calibrated phylogenies including more than 45 species (*74–76*).

Under the first approach, we calculated both distance- and frequency-based measures of convergence (*39*). Distance-based measures (C1–C4) are calculated between two lineages relative to their distance at the point in evolutionary history where the two lineages were maximally dissimilar. C1 specifically measures the proportion of phenotypic distance closed by evolution, ranging from 0 to 1 (where 1 indicates complete convergence). To calculate C1, ancestral states are reconstructed (via a Brownian motion model of evolution) for two or more putatively convergent lineages, back to their most recent common ancestor. The maximum phenotypic distance between any pair of ancestors (Dmax) is calculated, and compared with the phenotypic distance between the current putatively convergent taxa (Dtip). The greater the difference between Dmax and Dtip, the higher the index. C2 is the raw value of the difference between the maximum and extant distance between the two lineages. C3 is C2 scaled by the total evolution (sum of squared ancestor-to-descendant changes) between the two lineages. C4 is C2 scaled by the total evolution in the whole clade. These four measures quantify incomplete convergence in multidimensional space. Finally, C5, the frequency-based measure, quantifies and reports the number of convergent events where lineages evolve into a specific region of morphospace (crossing it from outside). C5 sums the number of times through the evolution of a clade that lineages evolve into a given region of phenotypic space. C5 is the number of focal taxa that reside within a limited but convergent region of a phylomorphospace (the phylogenetic connections between taxa represented graphically in a plot of morphological space). The significance of C1–C5 was found by running 1000 simulations for each comparison using Brownian-Motion on a variance–covariance matrix based on data-derived parameters, with convergence measures for each simulation calculated to determine if the observed C value is greater than expected by chance. A priori focal groups forming the basis of convergence tests were the same niche categorizations used in OUwie analyses. These analyses were performed using the R package *convevol* (*89*).

The second approach to measure convergence was based on comparing the angles formed by two tested clades from their most recent common ancestor with the expected angle according to null evolutionary models (*38*). Under the “state case”, *search.conv* computes the mean angle over all possible combinations of species pairs using one species per state. Each individual angle is divided by the patristic distance between the species. Significance is assessed by contrasting this value with a family of 1,000 random angles obtained by shuffling the state across the species (*38*). These analyses were performed using the R package *RRphylo* (*90*).

The third approach to measure convergence used the Wheatleaf metric (*40*). This index generates phenotypic (Euclidean) distances from any number of traits across species and penalizes them by phylogenetic distance before investigating similarity (in order to weight close phenotypic similarity higher for distantly related species). It uses an a priori designation of convergent species, which are defined as species belonging to a niche for which the traits are hypothesized to converge. The method then calculates a ratio of the mean (penalized) distances between all species to the mean (penalized) distances between allegedly convergent species. The index detects if convergent species diverge more in phenotypic space from the non-convergent species and show a tighter clustering to each other (*40*). The significance of this index was found by comparing the empirical values of the index with a distribution of simulated indices obtained running 5000 bootstrap simulations. These analyses were performed using the R package *windex* (*91*).

## Acknowledgments

Authors thank Raquel Sánchez, Angel Caravantes, Isabel Sánchez Almazo, María José Jorquera, and Iván Rodríguez Arós for helping us during several phases of the study. We also thank all contributors to the pollinator database (Table S10) for kindly sending us unpublished information on Brassicaceae floral visitors. This research is supported by grants from the Spanish Ministry of Science, Innovation and Universities (CGL2015-63827-P, CGL2017-86626-C2-1-P, CGL2017-86626-C2-2-P, UNGR15-CE-3315), Junta de Andalucía (P18-FR-3641, IE19_238 EEZA CSIC), LIFE18 GIE/IT/000755, and Xunta de Galicia (CITACA), including EU FEDER funds. This is a contribution to the Research Unit Modeling Nature, funded by the Consejería de Economía, Conocimiento, Empresas y Universidad, and European Regional Development Fund (ERDF), reference SOMM17/6109/UGR.

## SI FIGURES

**Figure S1.**
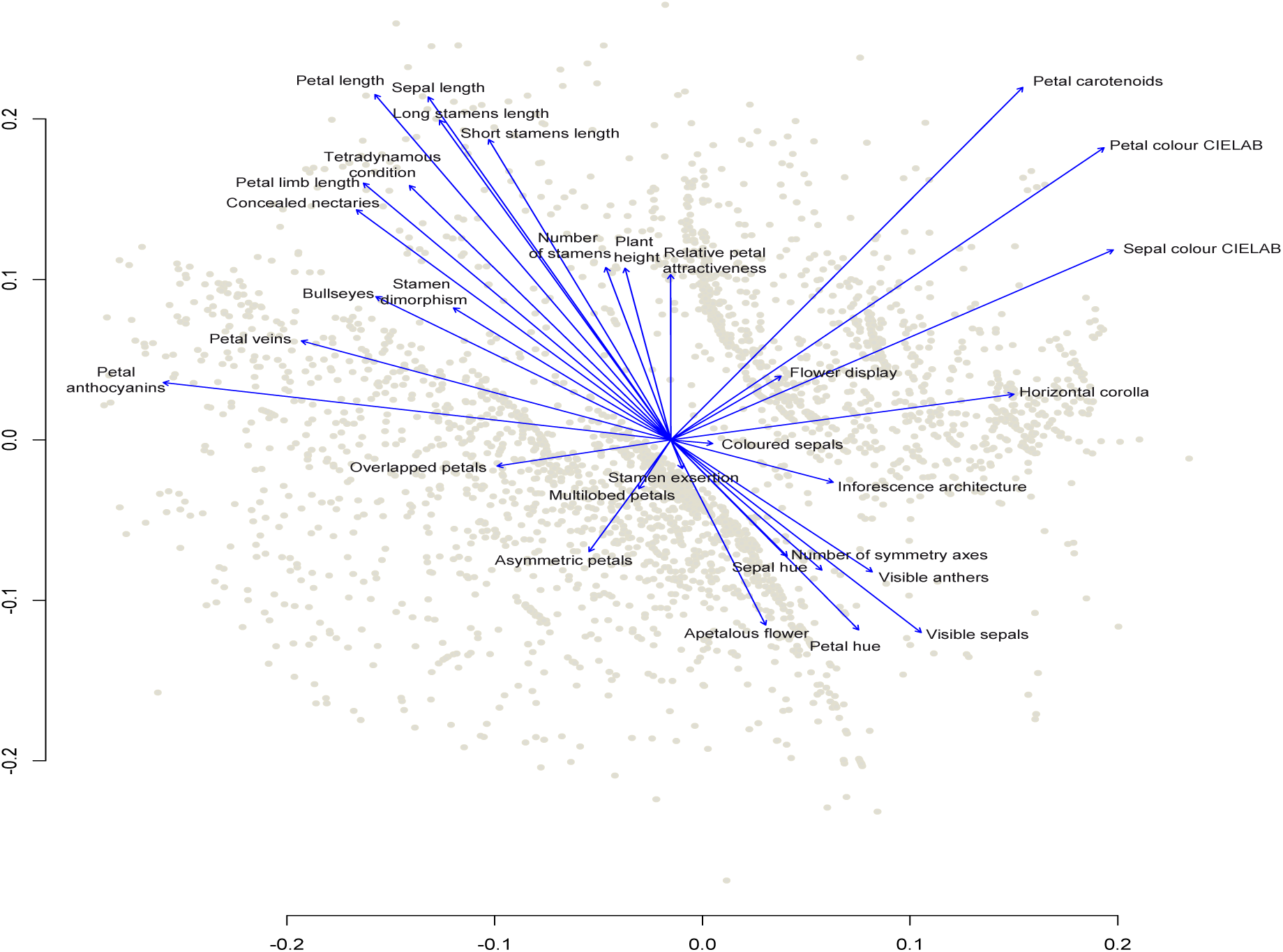
Association among the 31 pollination traits of 3140 Brassicaceae species. Trait vectors represent the Spearman correlations, with the length and direction indicating the relationship with composite NMDS axes.

**Table S1.**
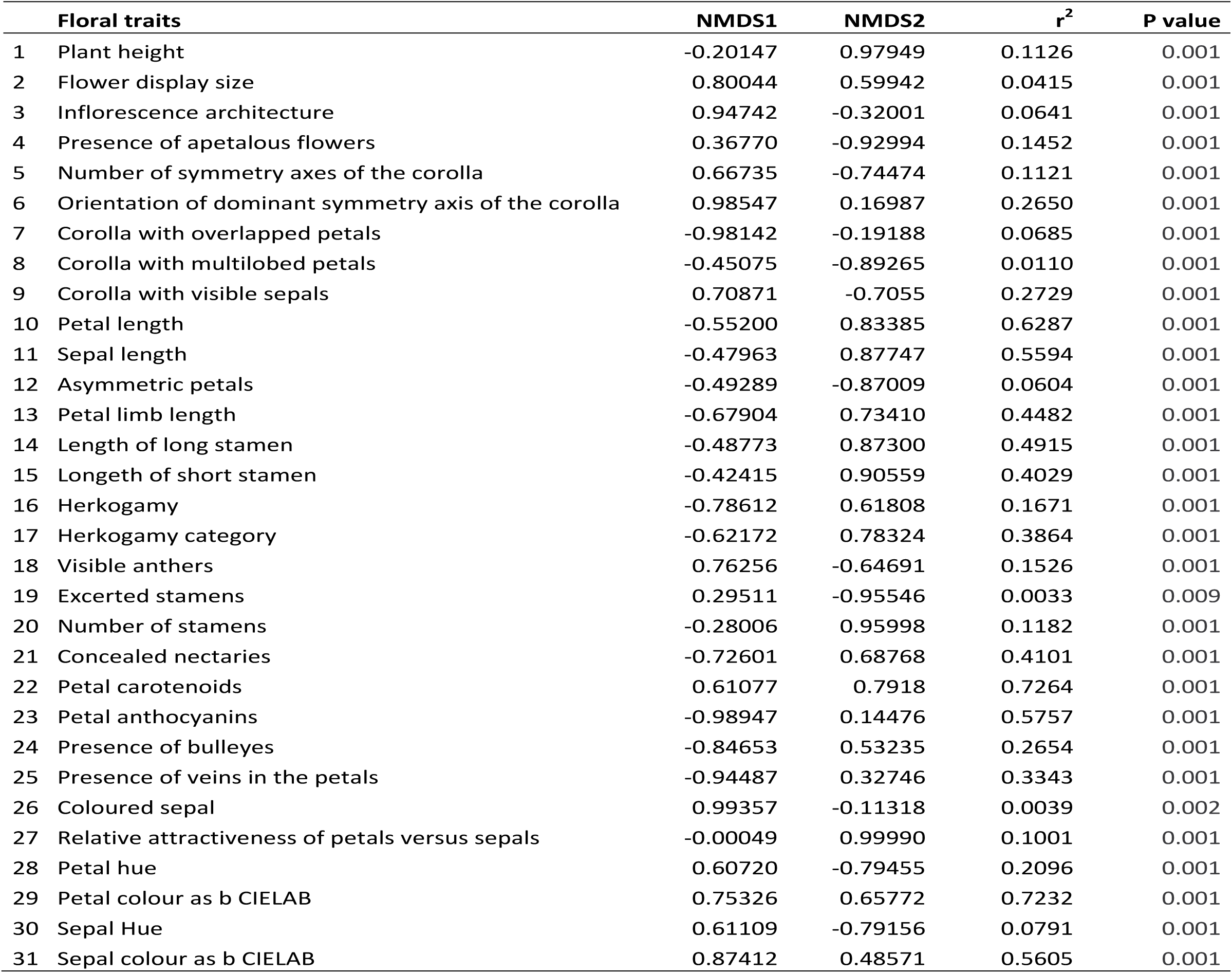
Fitting of the floral traits onto the NMDS vectors.

**Table S2.** Disparity, calculated as the Euclidean distance in the family-wide floral morphospace, between each of the 38 morphs included in our dataset (see Supplementary Data 2 for details and references) and their respective wild types.

| Type of polymorphism | Species | Morph | NMDS |
| --- | --- | --- | --- |
| Breeding system | *Brassica napus* | cleistogamous mutant | 0.035856865 |
| Breeding system | *Brassica rapa* | female sterility mutant | 0.133332180 |
| Breeding system | *Cardamine kokairensis* | cleistogamous mutant | 0.111696867 |
| Breeding system | *Leavenworthia crassa* | Outcrosser morph | 0.015771684 |
| Colour mutant | *Brassica napus* | white mutant | 0.173001555 |
| Colour mutant | *Moricandia arvensis* | white mutant | 0.149565069 |
| Flower colour polymorphism | *Boechnera stricta* | pink morph | 0.060580070 |
| Flower colour polymorphism | *Boechnera stricta* | purple morph | 0.054429426 |
| Flower colour polymorphism | *Cakile maritima* | white morph | 0.016547552 |
| Flower colour polymorphism | *Eruca vesicaria* | white morph | 0.082257145 |
| Flower colour polymorphism | *Erysimum cheiri* | purple cultivar | 0.073997237 |
| Flower colour polymorphism | *Erysimum cheiri* | white cultivar | 0.026460490 |
| Flower colour polymorphism | *Hesperis laciniata* | white morph | 0.069557518 |
| Flower colour polymorphism | *Hesperis matronalis* | white morph | 0.094873544 |
| Flower colour polymorphism | *Hormathophylla spinosa* | white morph | 0.101607789 |
| Flower colour polymorphism | *Leavenworthia stylosa* | white morph | 0.066269050 |
| Flower colour polymorphism | *Lobularia maritima* | deep purple cultivar | 0.090407642 |
| Flower colour polymorphism | *Marcus-kochia littorea* | light pink morph | 0.019542946 |
| Flower colour polymorphism | *Matthiola fruticulosa* | greenish morph | 0.008349920 |
| Flower colour polymorphism | *Matthiola incana* | white cultivar | 0.138357728 |
| Flower colour polymorphism | *Matthiola lunata* | white morph | 0.079964493 |
| Flower colour polymorphism | *Parrya nudicaulis* | white morph | 0.115226060 |
| Flower colour polymorphism | *Raphanus raphanistrum* | white morph | 0.082072459 |
| Flower colour polymorphism | *Raphanus raphanistrum* | yellow morph | 0.032685632 |
| Flower colour polymorphism | *Raphanus raphanistrum* | pink morph | 0.082778860 |
| Flower colour polymorphism | *Streptanthus glandulosus* | white morph | 0.014907493 |
| Gender dimorphism | *Hormathophylla spinosa* | female morph | 0.035783402 |
| Gender dimorphism | *Hormathophylla spinosa* | male morph | 0.025226337 |
| Gender dimorphism | *Lepidium sisymbrioides* | female morph | 0.067913758 |
| Gender dimorphism | *Lepidium solandri* | female morph | 0.065685257 |
| Gender dimorphism | *Pachycladon stellatum* | female morph | 0.014244744 |
| Gender dimorphism | *Pachycladon wallii* | male morph | 0.059320935 |
| Homeotic mutant | *Arabidopsis thaliana* | AGAMOUS mutant | 0.001052532 |
| Homeotic mutant | *Arabidopsis thaliana* | APETALA1 mutant | 0.119617731 |
| Homeotic mutant | *Arabidopsis thaliana* | APETALA3 mutant | 0.102445400 |
| Homeotic mutant | *Capsella bursapastoris* | Spe mutant | 0.051187659 |
| Phenotypic plasticity | *Cardamine hirsuta* | plastic change in stamens number | 0.010624208 |
| Phenotypic plasticity | *Cardamine hirsuta* | plastic change in petal number | 0.060755859 |

**Table S3.**
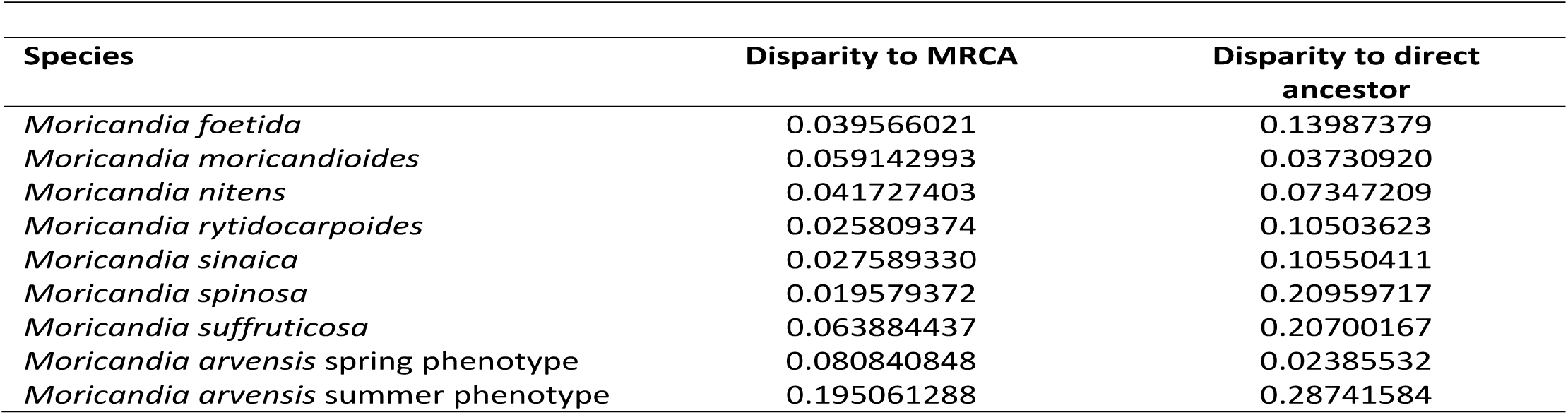
Floral disparity of each species of *Moricandia* from the most recent common ancestor (MRCA) of the genus and from the direct ancestor of each species.

| Species | Disparity to MRCA | Disparity to direct ancestor |
| --- | --- | --- |
| *Moricandia foetida* | 0.039566021 | 0.13987379 |
| *Moricandia moricandioides* | 0.059142993 | 0.03730920 |
| *Moricandia nitens* | 0.041727403 | 0.07347209 |
| *Moricandia rytidocarpoides* | 0.025809374 | 0.10503623 |
| *Moricandia sinaica* | 0.027589330 | 0.10550411 |
| *Moricandia spinosa* | 0.019579372 | 0.20959717 |
| *Moricandia suffruticosa* | 0.063884437 | 0.20700167 |
| *Moricandia arvensis* spring phenotype | 0.080840848 | 0.02385532 |
| *Moricandia arvensis* summer phenotype | 0.195061288 | 0.28741584 |

**Table S4.** Significance of the Mantel tests checking for spatial autocorrelation across the morphospace of the pollinator functional groups. Due to the small abundance of some pollinators, the original 43 functional groups have been pooled in 26 main functional groups.

| Functional Groups | Mantel R | p value |
| --- | --- | --- |
| Ant | 0.047 | 0.055 |
| Autogamy | 0.257 | 0.001 |
| Bug | 0.032 | 0.192 |
| Butterfly | 0.089 | 0.001 |
| Hawkmoth | 0.054 | 0.035 |
| Hoverfly | 0.072 | 0.003 |
| Large beefly | 0.043 | 0.079 |
| Large beetle | 0.046 | 0.063 |
| Large fly | 0.052 | 0.044 |
| Large wasp | 0.053 | 0.048 |
| Long tongued fly | 0.092 | 0.001 |
| Long tongued large bee | 0.242 | 0.001 |
| Long tongued medium-sized bee | 0.051 | 0.041 |
| Moth | 0.039 | 0.113 |
| Nocturnal moth | 0.222 | 0.001 |
| Other | 0.018 | 0.497 |
| Pollen wasp | 0.026 | 0.313 |
| Small beetle | 0.012 | 0.613 |
| Small diving beetle | 0.003 | 0.899 |
| Small fly | 0.112 | 0.001 |
| Small wasp | 0.041 | 0.112 |
| Short tongued large bee | 0.065 | 0.004 |
| Short tongued medium-sized bee | 0.012 | 0.590 |
| Short tongued small bee | 0.073 | 0.001 |
| Short tongued extra small bee | 0.020 | 0.454 |
| Thrips | -0.014 | 0.758 |

**Table S5.** Diferences between the two *Moricandia arvensis* phenotypes in the visitation frequency (both in absolute number of insects and in proportion of visits) of every pollinator functional group. Fifteen censuses of 1 hr and two researchers per phenotype.

| Pollinator functional group | spring phenotype | summer phenotype | spring phenotype (proportion) | summer phenotype (proportion) |
| --- | --- | --- | --- | --- |
| Hawkmoth | 91 | 0 | 0.036 | 0.000 |
| Honeybee | 40 | 0 | 0.016 | 0.000 |
| Large beefly | 309 | 72 | 0.124 | 0.148 |
| Large beetle | 30 | 40 | 0.012 | 0.082 |
| Large butterfly | 280 | 66 | 0.112 | 0.135 |
| Large fly | 38 | 0 | 0.015 | 0.000 |
| Large hoverfly | 8 | 7 | 0.003 | 0.014 |
| Long tongued large bee | 1131 | 5 | 0.453 | 0.010 |
| Long tongued medium-sized bee | 226 | 0 | 0.090 | 0.000 |
| Small beefly | 6 | 0 | 0.002 | 0.000 |
| Small beetle | 21 | 8 | 0.008 | 0.016 |
| Small butterfly | 11 | 0 | 0.004 | 0.000 |
| Small diving beetle | 25 | 12 | 0.010 | 0.025 |
| Small fly | 12 | 0 | 0.005 | 0.000 |
| Small hoverfly | 49 | 37 | 0.020 | 0.076 |
| Small moth | 2 | 7 | 0.001 | 0.014 |
| Short tongued large bee | 89 | 3 | 0.036 | 0.006 |
| Short tongued medium-sized bee | 36 | 0 | 0.014 | 0.000 |
| Short tongued small bee | 78 | 207 | 0.031 | 0.424 |
| Short tongued extra small bee | 1 | 0 | 0.000 | 0.000 |
| Thrips | 15 | 24 | 0.006 | 0.049 |

**Table S6.** Outcome of the analyses to test the occurrence of floral convergence among plants from niches 3 and 5. **Angle** is the mean theta angle between all species belonging to the same niche. **Angle/time** is the angle divided by time distance. The significance of these angles has been found by comparing with a null model consisting in shuffling each niche 1,000 times across the tree tips and calculating a distribution of random angle. **C1** measures the proportion of phenotypic distance closed by evolution, ranging from 0 to 1 (where 1 indicates complete convergence). **C2** is the raw value of the difference between the maximum and extant distance between the lineages. **C3** is C2 scaled by the total evolution (sum of squared ancestor-to-descendant changes) between the two lineages. **C4** is C2 scaled by the total evolution in the whole clade. The significance of C1-C2, was evaluated by running 1000 simulations for each comparison using Brownian-Motion models. **Wheatleaf** is the ratio of the mean (penalized) distances between all species to the mean (penalized) distances between allegedly convergent species. Significance found by running 2000 bootstrapping simulations. In bold, significant values.

| Phylogenies | Smith & Brown 2018 |  | Gaynor et al. 2018 |  | Huang et al. 2019 |  |
| --- | --- | --- | --- | --- | --- | --- |
|  | Value | p | Value | p | Value | p |
| **Niche 3** |  |  |  |  |  |  |
| Angle | **80.587** | 0.008 | **79.431** | 0.002 | **64.930** | 0.055 |
| Angle/time | 2.350 | 0.719 | 1.645 | 0.397 | 4.023 | 0.815 |
| C1 | **0.373** | 0.000 | **0.472** | 0.000 | **0.415** | 0.000 |
| C2 | **0.104** | 0.000 | **0.142** | 0.000 | **0.104** | 0.000 |
| C3 | **0.141** | 0.000 | **0.166** | 0.000 | **0.219** | 0.000 |
| C4 | 0.003 | 0.720 | 0.002 | 0.700 | 0.008 | 0.600 |
| Wheatleaf | 0.830 | 0.986 | 0.940 | 0.715 | **1.060** | 0.028 |
| **Niche 5** |  |  |  |  |  |  |
| Angle | **70.093** | 0.002 | **73.491** | 0.002 | **58.313** | 0.049 |
| Angle/time | **1.393** | 0.021 | 1.783 | 0.745 | **2.474** | 0.011 |
| C1 | **0.356** | 0.000 | **0.472** | 0.000 | **0.240** | 0.000 |
| C2 | **0.110** | 0.000 | **0.142** | 0.000 | **0.075** | 0.000 |
| C3 | **0.128** | 0.000 | **0.166** | 0.000 | **0.118** | 0.000 |
| C4 | 0.003 | 0.727 | 0.002 | 0.700 | 0.006 | 0.545 |
| Wheatleaf | 1.120 | 0.673 | **1.170** | 0.094 | 0.920 | 0.978 |

**Table S7.** Outcome of the analyses testing for morphological convergence between the *Moricandia* clade and the rest of clades included in each time-calibrated phylogeny. **Clade size** is the number of species within the *Moricandia* clade. **θ_real_** is the mean angle over all possible combinations of pairs of species taking one species per clade. **θ_ace_** is the mean angle between ancestral states between each pairs of clades. **dist_mrca_** is the patristic distance (sum of brach length) between the most recent common ancestors of each pair of clade. We indicate the congervent clades and the pollination niches of each species included in the convergent clades. In red *Moricandia* clades including *Moricandia arvensis* spring phenotype. Tribes (E= Erysimeae, A= Anchonieae, C=Cardamineae, M=Malcolmieae, An=Anastaticeae).

| Moricandia clade | Clade 2 | Clade size | $\theta_{\text{real}}$ | $\theta_{\text{ace}}$ | $\text{dist}_{\text{mrca}}$ | $\theta_{\text{real}}/\text{dist}_{\text{mrca}}$ | $\theta_{\text{real}}/\text{dist}_{\text{mrca}}$ p-value | $\theta_{\text{ace}}+\theta_{\text{real}}/\text{dist}_{\text{mrca}}$ | $\theta_{\text{ace}}+\theta_{\text{real}}/\text{dist}_{\text{mrca}}$ p-value | Convergent clades | Tribe | Niche |
| --- | --- | --- | --- | --- | --- | --- | --- | --- | --- | --- | --- | --- |
| **Smith & Brown's phylogeny** |  |  |  |  |  |  |  |  |  |  |  |  |
| 253 | 347 | 7 | 15.200 | 4.420 | 124.236 | 0.122 | 0.058 | 0.158 | 0.012 | *Erysimum bicolor/ scoparium* | E | 5,5 |
| 254 | 347 | 5 | 19.035 | 5.389 | 129.718 | 0.147 | 0.058 | 0.188 | 0.016 | *Erysimum bicolor/ scoparium* | E | 5,5 |
| 255 | 347 | 4 | 22.601 | 6.331 | 133.342 | 0.169 | 0.052 | 0.217 | 0.019 | *Erysimum bicolor/ scoparium* | E | 5,5 |
| 256 | 347 | 2 | 6.180 | 7.281 | 133.538 | 0.046 | 0.026 | 0.101 | 0.005 | *Erysimum bicolor/ scoparium* | E | 5,5 |
| 256 | 316 | 2 | 12.950 | 2.109 | 59.810 | 0.217 | 0.061 | 0.252 | 0.018 | *Matthiola* clade | A | 6,1 |
| 256 | 375 | 2 | 8.543 | 13.786 | 49.285 | 0.173 | 0.066 | 0.453 | 0.045 | *Cardamine pentaphyllo/ pratensis* | C | 3,7 |
| 256 | 405 | 2 | 8.616 | 1.048 | 44.577 | 0.193 | 0.068 | 0.217 | 0.022 | *Malcolmia maritima— Marcus-kochia ramosissima* | M/An | 5,7 |
| 256 | 335 | 2 | 28.715 | 33.529 | 128.518 | 0.223 | 0.077 | 0.484 | 0.049 | *Erysimum popovii/ bastetanum/ semperflorens* | E | 5,5,6 |
| 257 | 347 | 2 | 39.021 | 6.003 | 133.462 | 0.292 | 0.103 | 0.337 | 0.015 | *Erysimum bicolor/ scoparium* | E | 5,5 |
| 258 | 347 | 2 | 5.613 | 4.410 | 124.383 | 0.045 | 0.012 | 0.081 | 0.002 | *Erysimum bicolor/ scoparium* | E | 5,5 |
| **Gaynor et al.'s phylogeny** |  |  |  |  |  |  |  |  |  |  |  |  |
| 481 | 334 | 2 | 10.943 | 17.832 | 78.194 | 0.140 | 0.052 | 0.368 | 0.037 | *Erysimum bicolor/ scoparium* | E | 5,5 |
| 479 | 334 | 2 | 9.357 | 12.654 | 81.141 | 0.115 | 0.052 | 0.271 | 0.033 | *Erysimum bicolor/ scoparium* | E | 5,5 |
| 479 | 333 | 2 | 12.809 | 14.978 | 81.151 | 0.158 | 0.070 | 0.342 | 0.049 | *Erysimum lagascae/ rondae* | E | 5,3 |
| **Huang et al.'s phylogeny** |  |  |  |  |  |  |  |  |  |  |  |  |
| 143 | 83 | 2 | 4.960 | 0.985 | 23.966 | 0.207 | 0.023 | 0.248 | 0.001 | *Erucaria* clade | B | 3,5 |
| 143 | 82 | 2 | 7.094 | 0.554 | 22.049 | 0.322 | 0.041 | 0.347 | 0.002 | *Erucaria* clade + *Cakile* clade | B | 3,3,3,5 |
| 143 | 81 | 2 | 6.651 | 40.317 | 18.008 | 0.369 | 0.046 | 2.608 | 0.170 | *Erucaria* clade + *Cakile* clade + *Eremophyton chevallieri* | B | 3,3,3,5,5 |
| 143 | 84 | 2 | 9.229 | 1.876 | 23.466 | 0.393 | 0.066 | 0.473 | 0.003 | *Cakile* clade | B | 3,3 |
| 143 | 86 | 2 | 11.634 | 12.691 | 24.281 | 0.479 | 0.074 | 1.002 | 0.026 | *Zilla* clade | B | 5,5 |
| 143 | 85 | 2 | 9.383 | 12.437 | 24.007 | 0.391 | 0.076 | 0.909 | 0.029 | *Zilla* clade + *Foleyola billotii* | B | 3,5,5 |

**Table S8.**
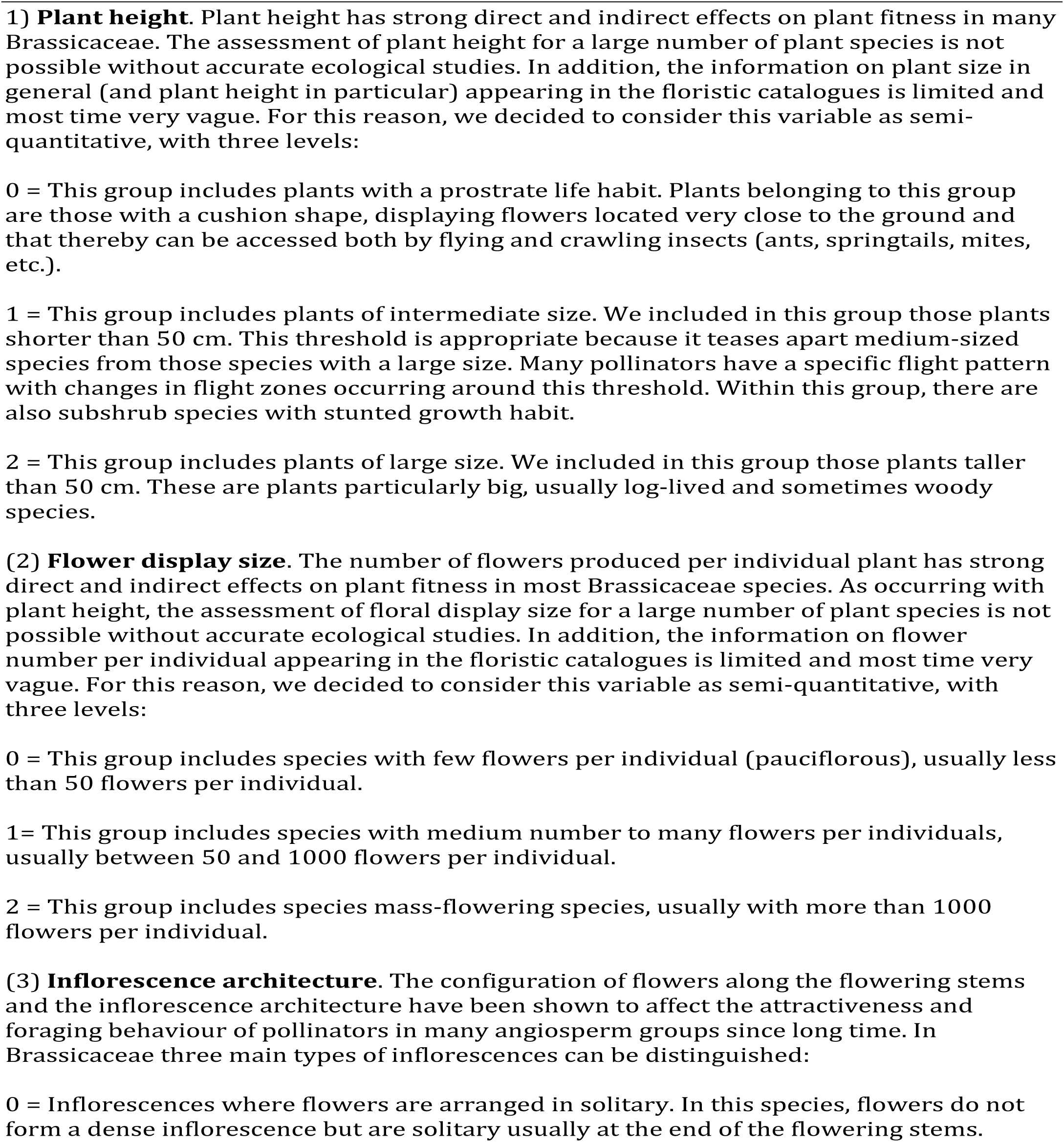
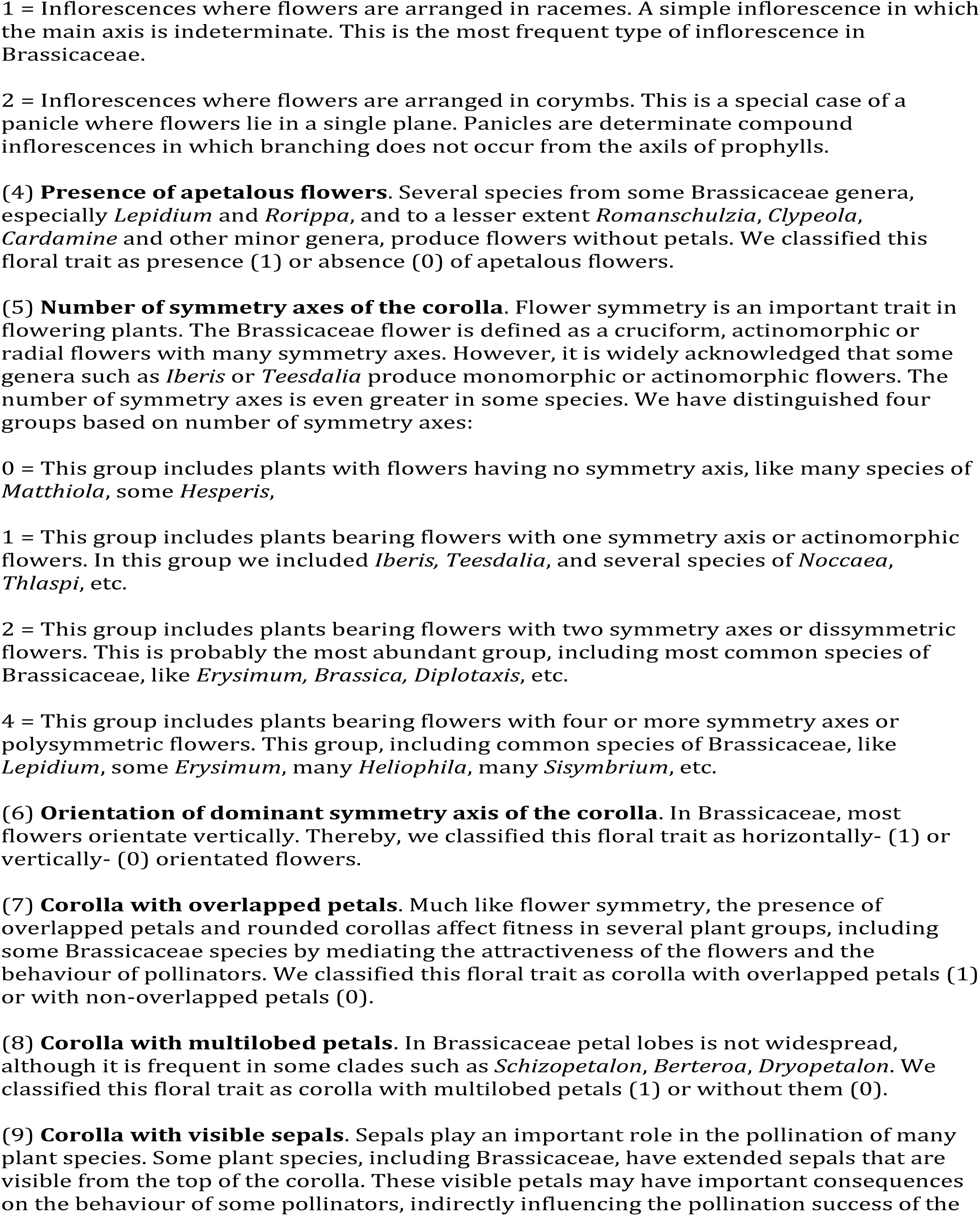
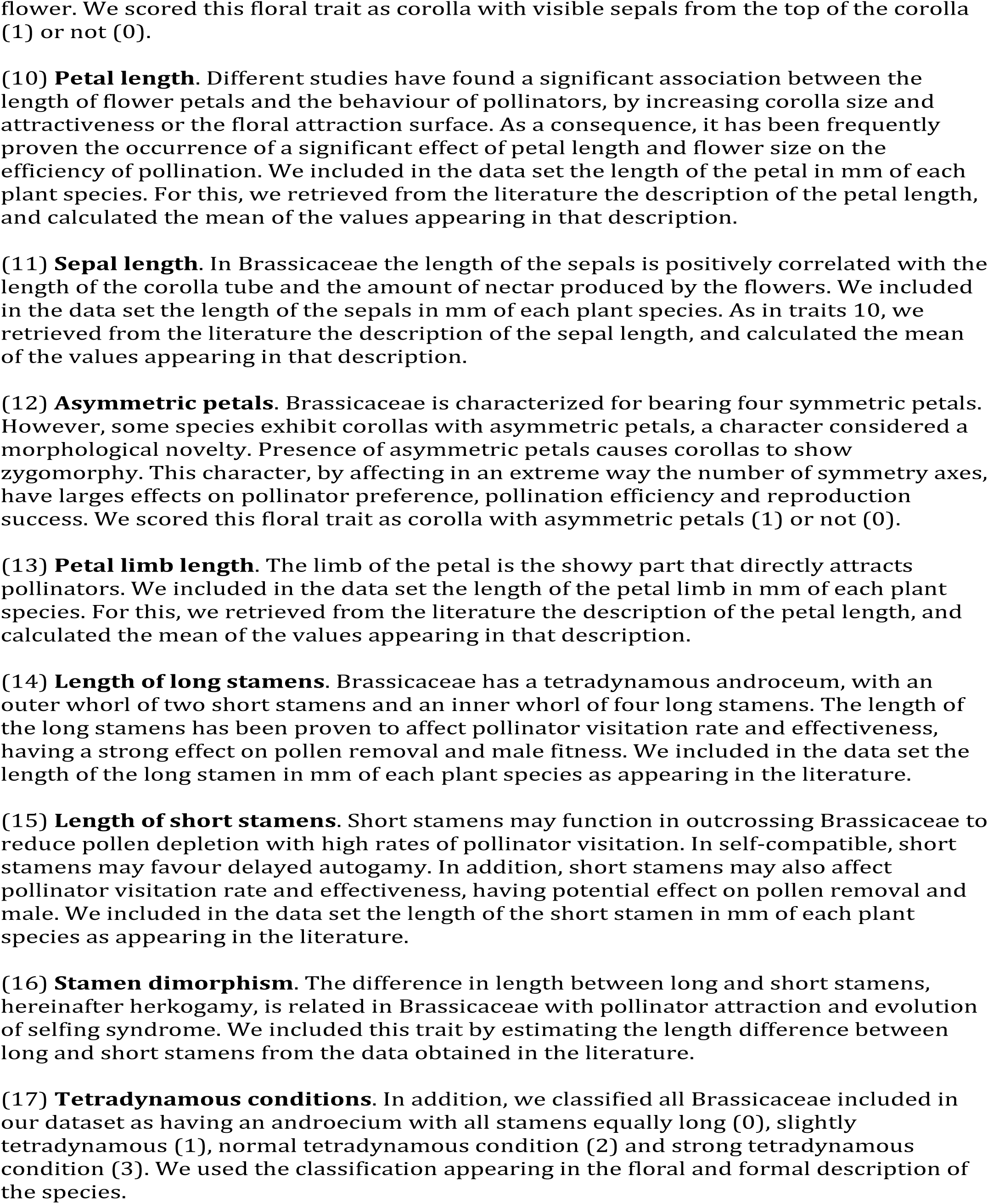
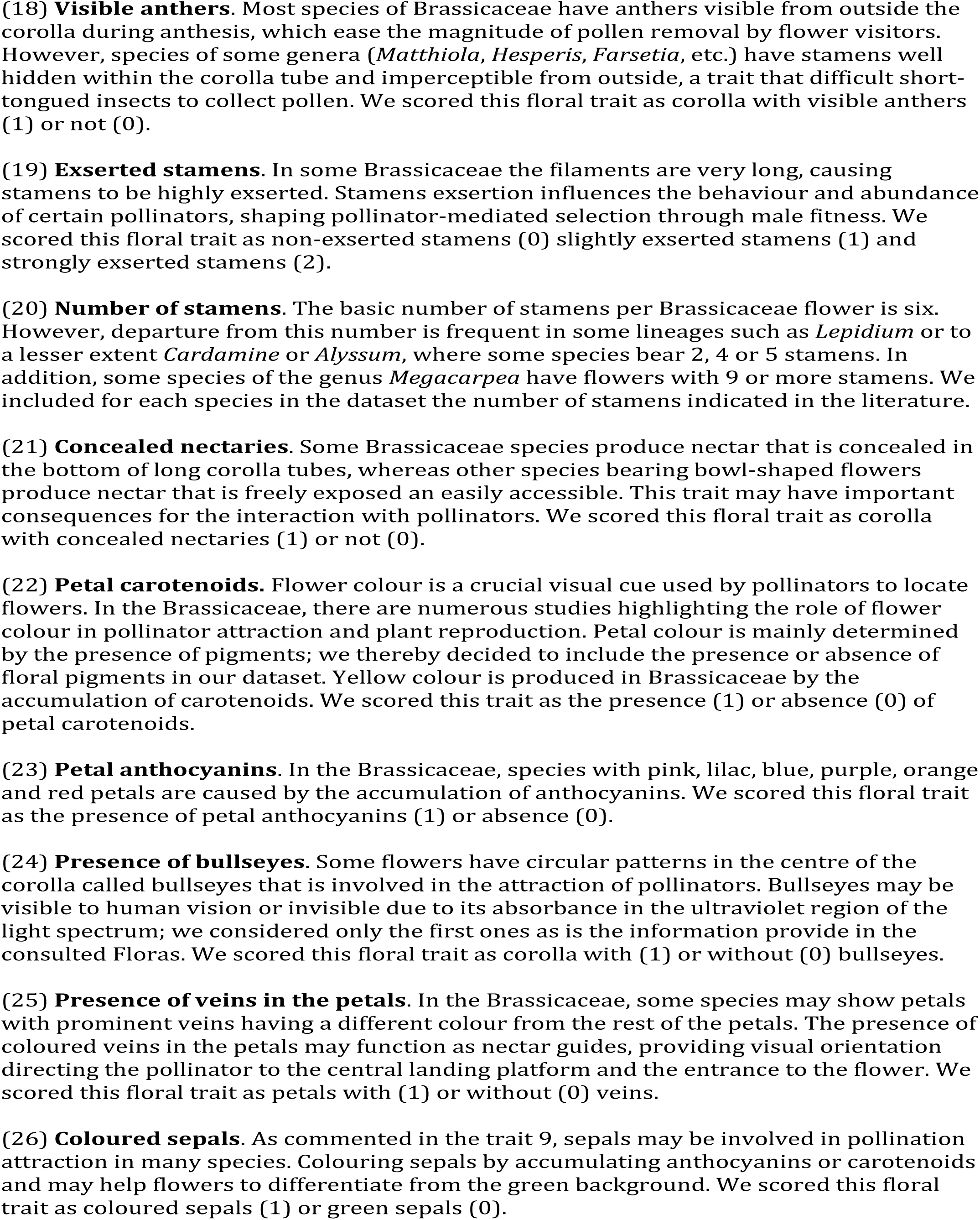
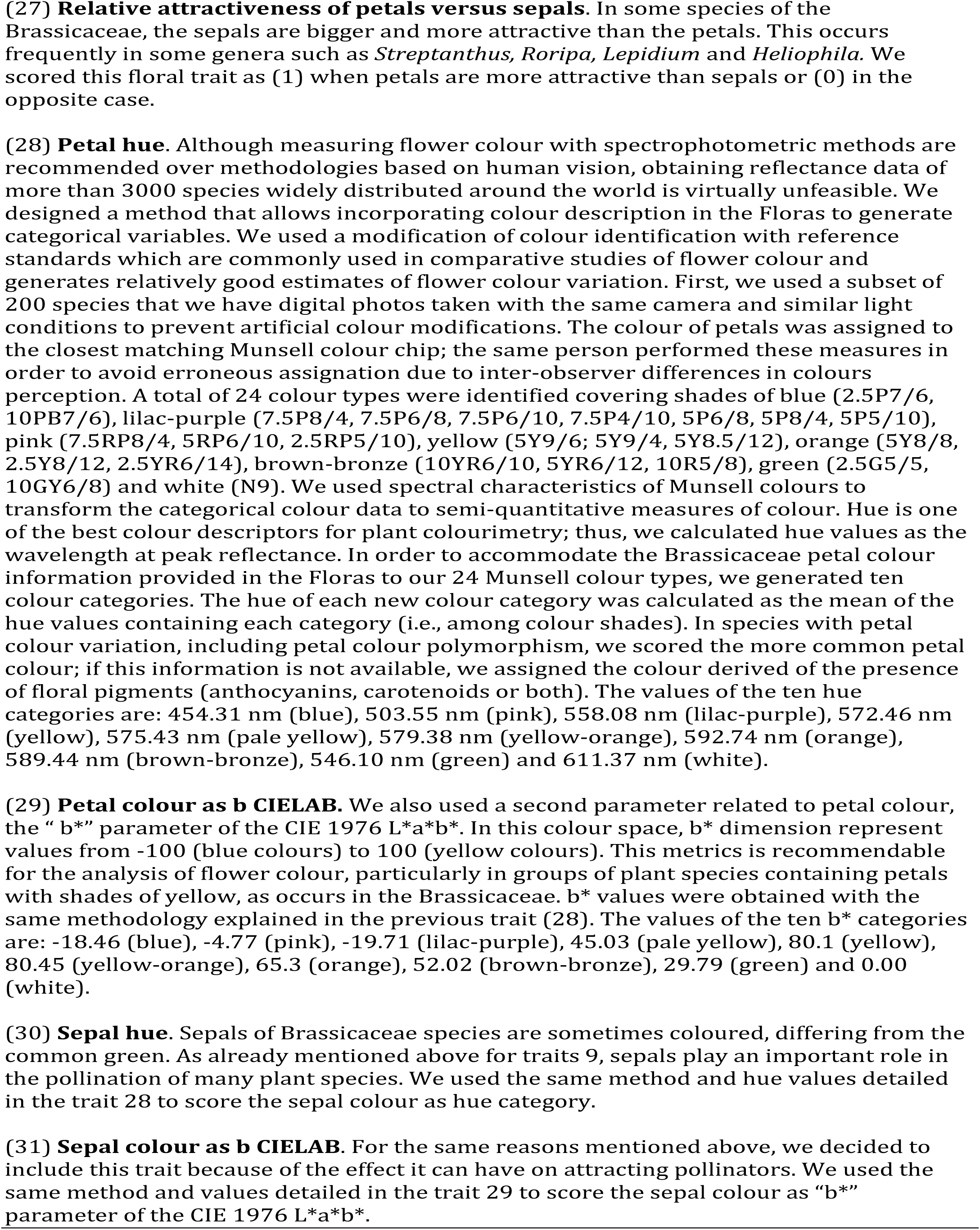
Description of floral traits related to pollinator attraction used to generate the floral morphospace in Brassicaceae. Pollinators respond to the variability of numerous phenotypic traits of plants, and the magnitude of their response shapes the reproductive success of the plants. We estimated for each plant included in our data set the values of several important floral traits.

**Table S9.**
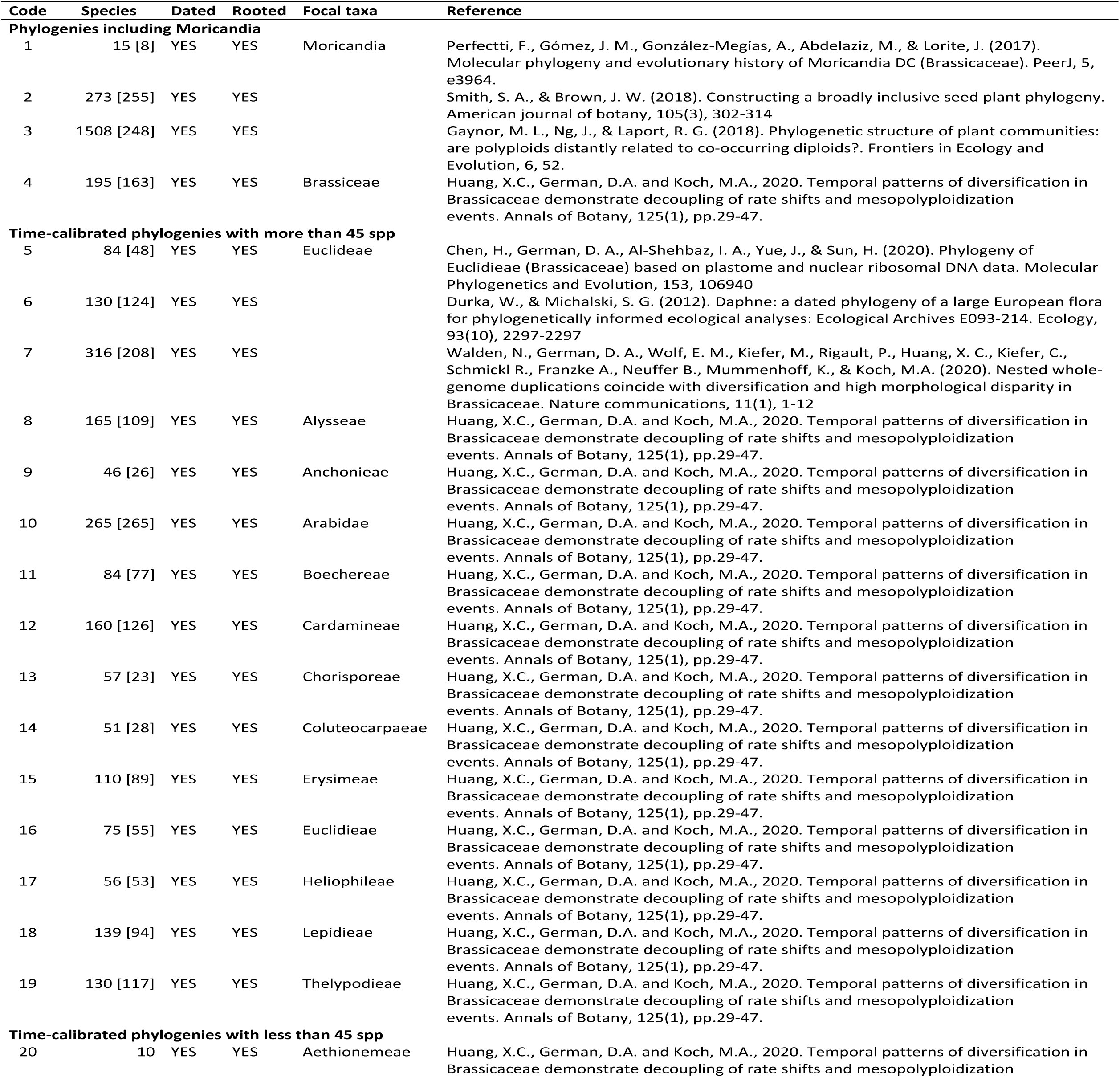
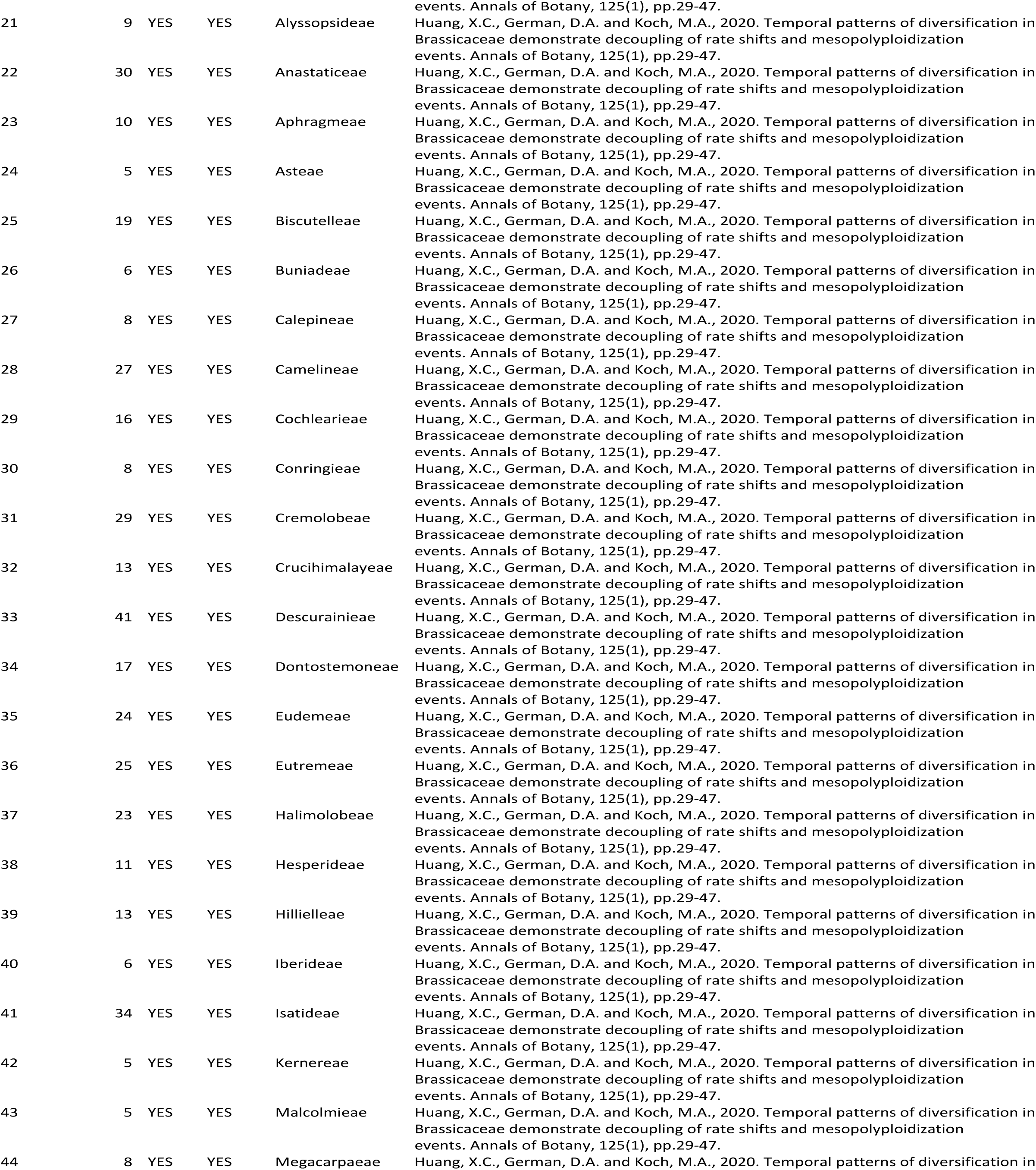
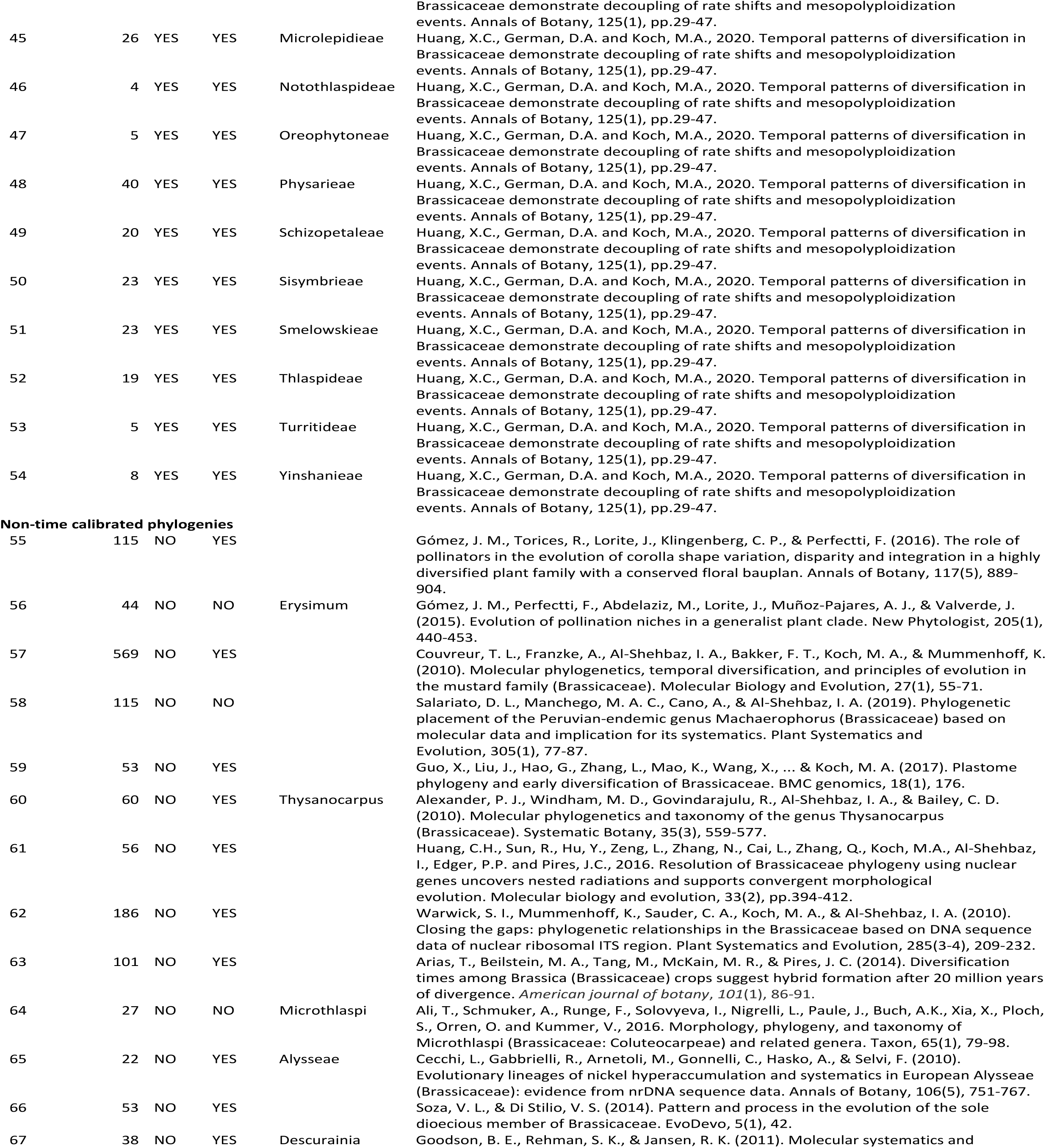
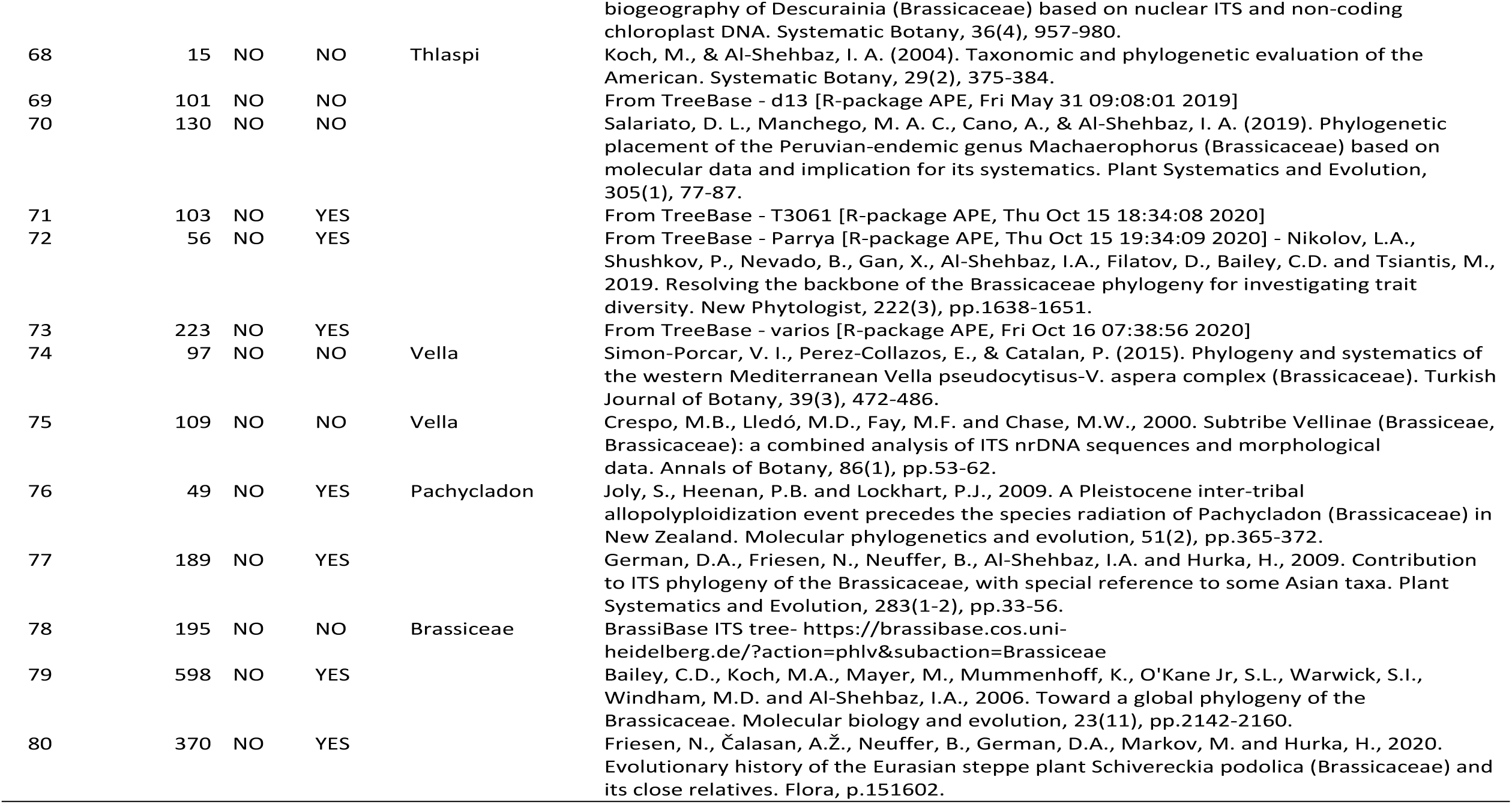
List of the phylogenies retrieved from the online repositories and from the literature to built up the Brassicaceae supertree. Within brackets appears the number of species included in the analysis of disparity

**Table S10.**
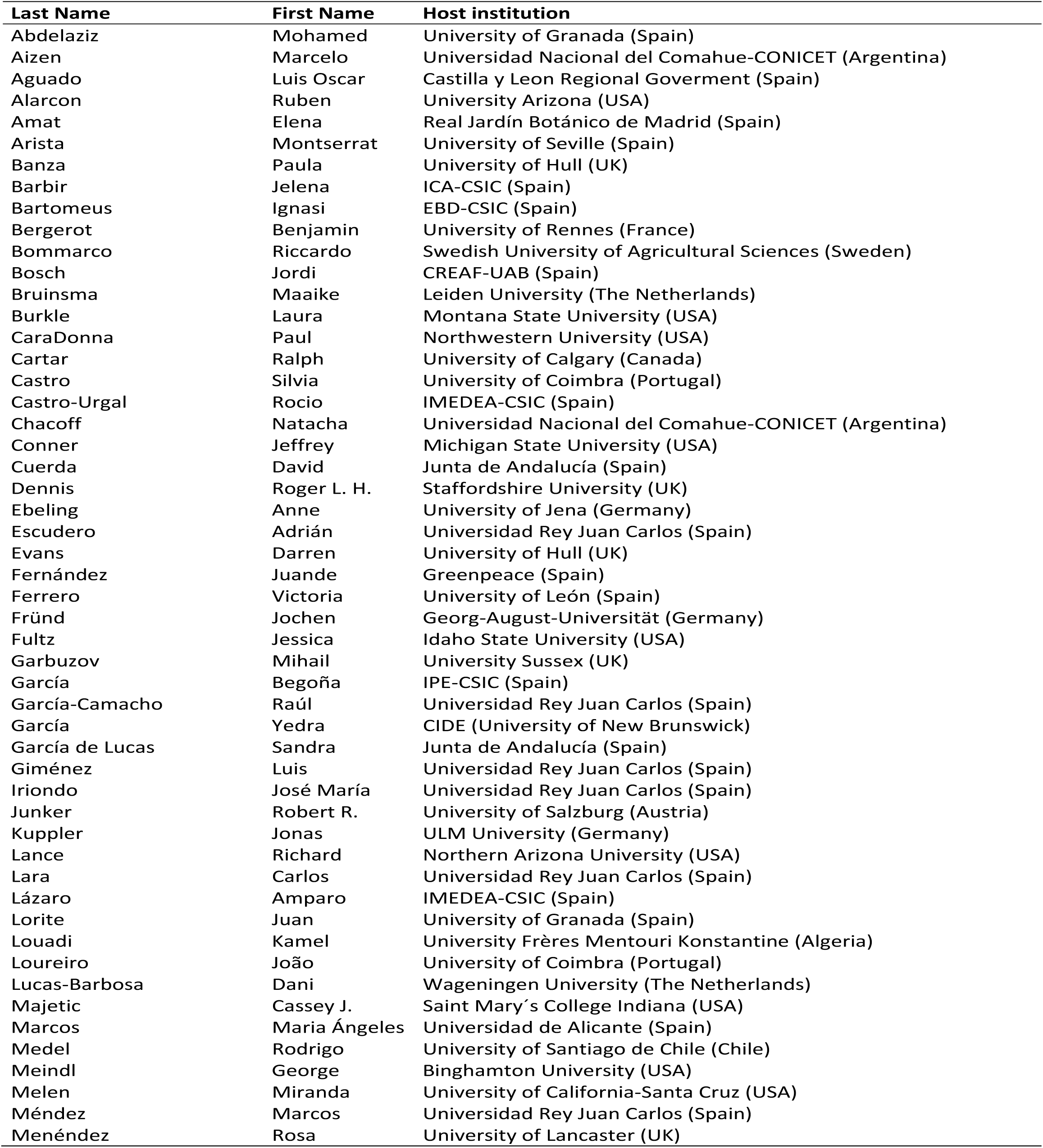
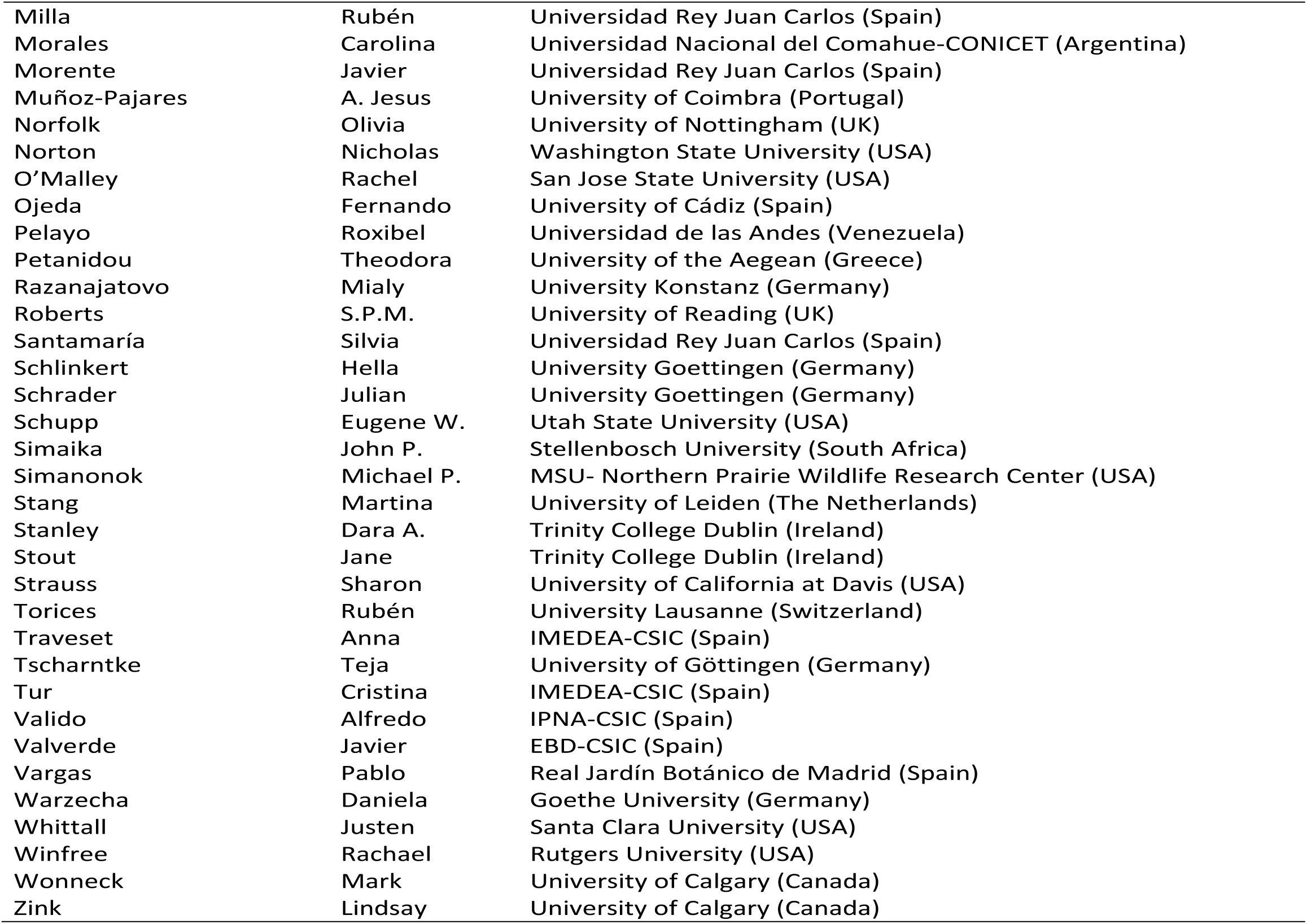
List of ecologists kindly sharing unpublished information on Brassicaceae pollinators. The host institutions are those at the time of the contact with our team.

| **Last Name** | **First Name** | **Host institution** |
| --- | --- | --- |
| Abdelaziz | Mohamed | University of Granada (Spain) |
| Aizen | Marcelo | Universidad Nacional del Comahue-CONICET (Argentina) |
| Aguado | Luis Oscar | Castilla y Leon Regional Government (Spain) |
| Alarcon | Ruben | University Arizona (USA) |
| Amat | Elena | Real Jardín Botánico de Madrid (Spain) |
| Arista | Montserrat | University of Seville (Spain) |
| Banza | Paula | University of Hull (UK) |
| Barbir | Jelena | ICA-CSIC (Spain) |
| Bartomeus | Ignasi | EBD-CSIC (Spain) |
| Bergerot | Benjamin | University of Rennes (France) |
| Bommarco | Riccardo | Swedish University of Agricultural Sciences (Sweden) |
| Bosch | Jordi | CREAF-UAB (Spain) |
| Bruinsma | Maaïke | Leiden University (The Netherlands) |
| Burkle | Laura | Montana State University (USA) |
| CaraDonna | Paul | Northwestern University (USA) |
| Cartar | Ralph | University of Calgary (Canada) |
| Castro | Silvia | University of Coimbra (Portugal) |
| Castro-Urgal | Rocio | IMEDEA-CSIC (Spain) |
| Chacoff | Natacha | Universidad Nacional del Comahue-CONICET (Argentina) |
| Conner | Jeffrey | Michigan State University (USA) |
| Cuerda | David | Junta de Andalucía (Spain) |
| Dennis | Roger L. H. | Staffordshire University (UK) |
| Ebeling | Anne | University of Jena (Germany) |
| Escudero | Adrián | Universidad Rey Juan Carlos (Spain) |
| Evans | Darren | University of Hull (UK) |
| Fernández | Juande | Greenpeace (Spain) |
| Ferrero | Victoria | University of León (Spain) |
| Fründ | Jochen | Georg-August-Universität (Germany) |
| Fultz | Jessica | Idaho State University (USA) |
| Garbuzov | Mihail | University Sussex (UK) |
| García | Begoña | IPE-CSIC (Spain) |
| García-Camacho | Raúl | Universidad Rey Juan Carlos (Spain) |
| García | Yedra | CIDE (University of New Brunswick) |
| García de Lucas | Sandra | Junta de Andalucía (Spain) |
| Giménez | Luis | Universidad Rey Juan Carlos (Spain) |
| Iriondo | José María | Universidad Rey Juan Carlos (Spain) |
| Junker | Robert R. | University of Salzburg (Austria) |
| Kuppler | Jonas | ULM University (Germany) |
| Lance | Richard | Northern Arizona University (USA) |
| Lara | Carlos | Universidad Rey Juan Carlos (Spain) |
| Lázaro | Amparo | IMEDEA-CSIC (Spain) |
| Lorite | Juan | University of Granada (Spain) |
| Louadi | Kamel | University Frères Mentouri Konstantine (Algeria) |
| Loureiro | João | University of Coimbra (Portugal) |
| Lucas-Barbosa | Dani | Wageningen University (The Netherlands) |
| Majetic | Cassey J. | Saint Mary's College Indiana (USA) |
| Marcos | Maria Ángeles | Universidad de Alicante (Spain) |
| Medel | Rodrigo | University of Santiago de Chile (Chile) |
| Meindl | George | Binghamton University (USA) |
| Melen | Miranda | University of California-Santa Cruz (USA) |
| Méndez | Marcos | Universidad Rey Juan Carlos (Spain) |
| Menéndez | Rosa | University of Lancaster (UK) |
| Milla | Rubén | Universidad Rey Juan Carlos (Spain) |
| Morales | Carolina | Universidad Nacional del Comahue-CONICET (Argentina) |
| Morente | Javier | Universidad Rey Juan Carlos (Spain) |
| Muñoz-Pajares | A. Jesus | University of Coimbra (Portugal) |
| Norfolk | Olivia | University of Nottingham (UK) |
| Norton | Nicholas | Washington State University (USA) |
| O'Malley | Rachel | San Jose State University (USA) |
| Ojeda | Fernando | University of Cádiz (Spain) |
| Pelayo | Roxibel | Universidad de las Andes (Venezuela) |
| Petanidou | Theodora | University of the Aegean (Greece) |
| Razanajatovo | Mialy | University Konstanz (Germany) |
| Roberts | S.P.M. | University of Reading (UK) |
| Santamaría | Silvia | Universidad Rey Juan Carlos (Spain) |
| Schlinkert | Hella | University Goettingen (Germany) |
| Schrader | Julian | University Goettingen (Germany) |
| Schupp | Eugene W. | Utah State University (USA) |
| Simaika | John P. | Stellenbosch University (South Africa) |
| Simanonok | Michael P. | MSU- Northern Prairie Wildlife Research Center (USA) |
| Stang | Martina | University of Leiden (The Netherlands) |
| Stanley | Dara A. | Trinity College Dublin (Ireland) |
| Stout | Jane | Trinity College Dublin (Ireland) |
| Strauss | Sharon | University of California at Davis (USA) |
| Torices | Rubén | University Lausanne (Switzerland) |
| Traveset | Anna | IMEDEA-CSIC (Spain) |
| Tscharntke | Teja | University of Göttingen (Germany) |
| Tur | Cristina | IMEDEA-CSIC (Spain) |
| Valido | Alfredo | IPNA-CSIC (Spain) |
| Valverde | Javier | EBD-CSIC (Spain) |
| Vargas | Pablo | Real Jardín Botánico de Madrid (Spain) |
| Warzecha | Daniela | Goethe University (Germany) |
| Whittall | Justen | Santa Clara University (USA) |
| Winfree | Rachael | Rutgers University (USA) |
| Wonneck | Mark | University of Calgary (Canada) |
| Zink | Lindsay | University of Calgary (Canada) |

**Table S11.**
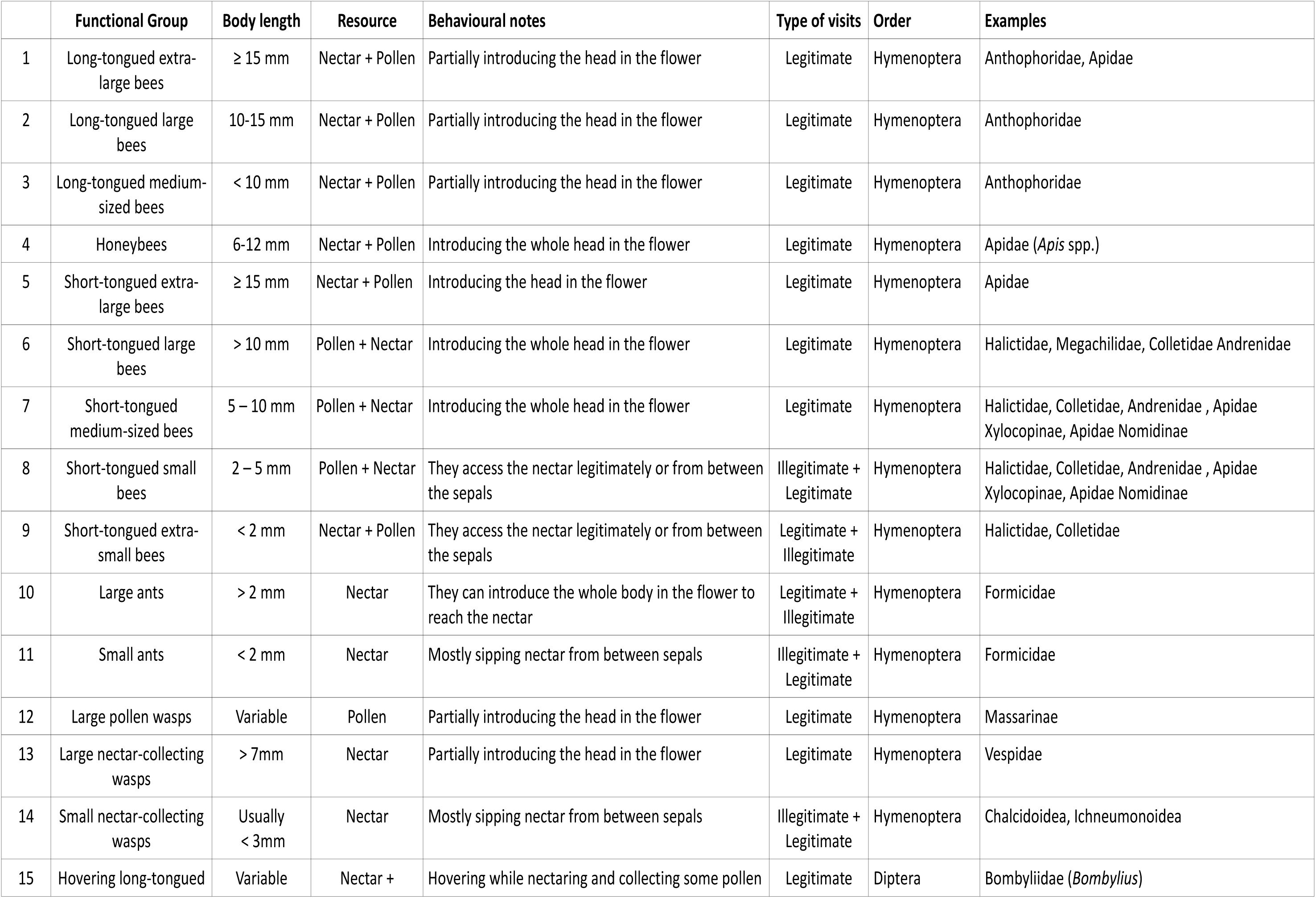
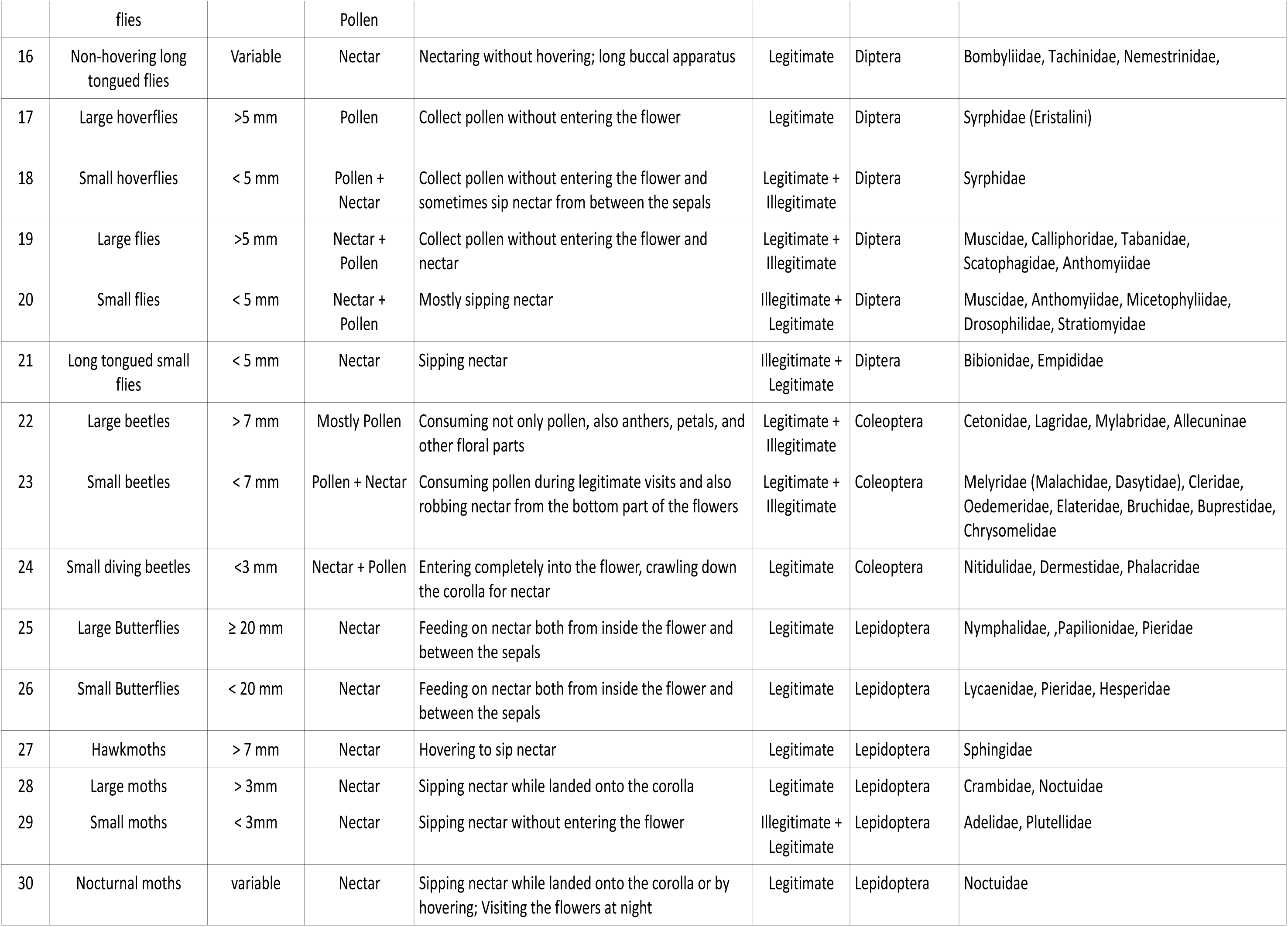
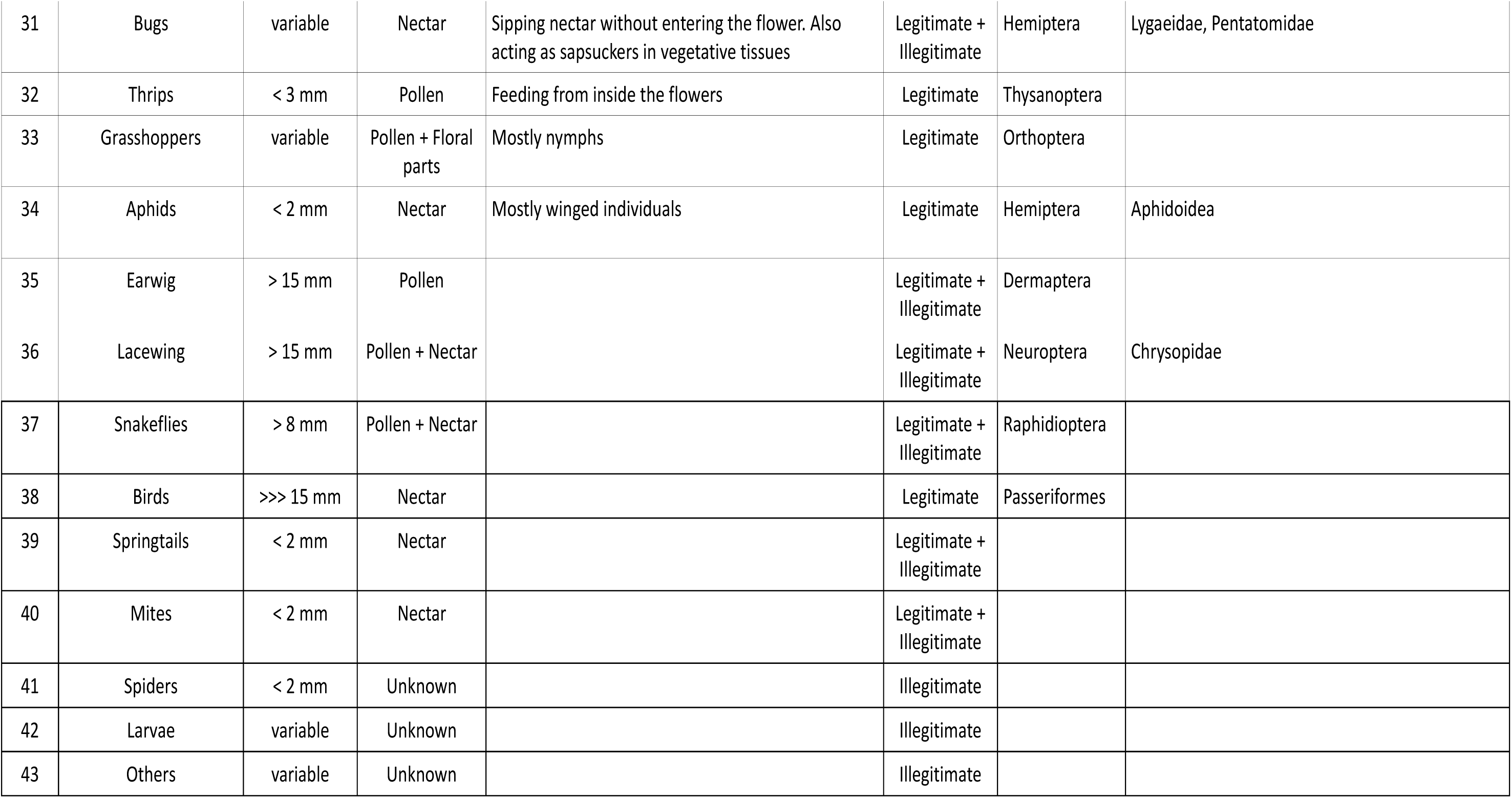
Brief description of the functional groups of the insects visiting the flowers of the studied species.

## References

1. G. R. McGhee, Convergent Evolution: Limited Forms Most Beautiful. MIT Press, Cambridge (2011). ISBN:9780262016421

2. J. B. Losos, Convergence, adaptation, and constraint. Evolution 65, 1827–1840 (2011). doi:10.1111/j.1558-5646.2011.01289.x

3. S. C. Morris, Life’s solution: Inevitable Humans in a Lonely Universe. Cambridge University Press, Cambridge (2003). ISBN:9780521603256

4. T. Pearce, Convergence and parallelism in evolution: a Neo-Gouldian account. Br. J. Philos. Sci. 63, 429–448 (2011). doi:10.1093/bjps/axr046

5. D. Schluter, The Ecology of Adaptive Radiation. Oxford Univ. Press, Oxford, U.K. (2000) ISBN:9780198505228

6. P. Nosil, Ecological Speciation. Oxford Univ. Press, Oxford, U.K. (2012) ISBN:9780199587100

7. N. Walden, D. A. German, E. M. Wolf, M. Kiefer, P. Rigault, X. C. Huang, C. Kiefer, R. Schmickl, A. Franzke, B. Neuffer, K. Mummenhoff, Nested whole-genome duplications coincide with diversification and high morphological disparity in Brassicaceae. Nature Comm. 11, 3795 (2020). doi:10.1038/s41467-020-17605-7

8. M. Simões, L. Breitkreuz, M. Alvarado, S. Baca, J. C. Cooper, L. Heins, K. Herzog, B. S. Lieberman, The evolving theory of evolutionary radiations. Trends Ecol. Evol. 31, 27–34 (2016). doi:10.1016/j.tree.2015.10.007

9. M. R., Pie, J. S. Weitz, A null model of morphospace occupation. Am. Nat. 166, E1–E13 (2005). doi:10.1086/430727

10. K. O. Winemiller, D. B. Fitzgerald, L. M. Bower, E. R. Pianka, Functional traits, convergent evolution, and periodic tables of niches. Ecol. Lett. 18, 737–751 (2015). doi:10.1111/ele.12462

11. C. T. Stayton, Are our phylomorphospace plots so terribly tangled? An investigation of disorder in data simulated under adaptive and nonadaptive models. Curr. Zool. 66, 565–574 (2020). doi:10.1093/cz/zoaa045

12. A. L. Pigot, C. Sheard, E. T. Miller, T. P. Bregman, B. G. Freeman, U. Roll, N. Seddon, C. H. Trisos, B. C. Weeks, J. A. Tobias, Macroevolutionary convergence connects morphological form to ecological function in birds. Nature Ecol. Evol. 4, 230–239 (2020). doi:10.1038/s41559-019-1070-4

13. D. L. Stern, The genetic causes of convergent evolution. Nat. Rev. Genet. 14, 751–764 (2013). doi:10.1038/nrg3483

14. E. B. Rosenblum, C. E. Parent, E. E. Brandt, The molecular basis of phenotypic convergence. Annu. Rev. Ecol. Evol. Syst. 45, 203–226 (2014). doi:10.1146/annurev-ecolsys-120213-091851

15. P. A. Christin, D. M. Weinreich, G. Besnard, Causes and evolutionary significance of genetic convergence. Trends Genet. 26, 400–405 (2010). doi:10.1016/j.tig.2010.06.005

16. M. J. West-Eberhard, Developmental Plasticity and Evolution. Oxford Univ. Press, New York, (2003). ISBN: 9780195122343

17. S. Sultan, Organism and Environment: Ecological development, Niche construction, and Adaption. Oxford Univ. Press, New York, (2015). ISBN:9780199587063

18. J. M. Gómez, F. Perfectti, C. Armas, E. Narbona, A. González-Megías, L. Navarro, L. DeSoto, R. Torices, Within-individual phenotypic plasticity in flowers fosters pollination niche shift. Nature Comm. 11, 4019 (2020). doi:10.1038/s41467-020-17875-1

19. V. Susoy, E. J. Ragsdale, N. Kanzaki, R. J. Sommer, Rapid diversification associated with a macroevolutionary pulse of developmental plasticity, eLife 4, e05463 (2015). doi:10.7554/eLife.05463

20. N. A. Levis, D. W. Pfennig, Evaluating ‘plasticity-first’ evolution in nature: key criteria and empirical approaches. Trends Ecol. Evol. 31, 563–574 (2016). doi:10.1016/j.tree.2016.03.012

21. R. J. Sommer, Phenotypic plasticity: from theory and genetics to current and future challenges. Genetics 215, 1–13 (2020). doi:10.1534/genetics.120.303163

22. M. A. Koch, D. A. German, M. Kiefer, A. Franzke, Database taxonomics as key to modern plant biology. Trends Plant Sci. 23, 4–6 (2018). doi:10.1016/j.tplants.2017.10.005

23. M. Kiefer, R. Schmickl, D. A. German, T. Mandáková, M. A. Lysak, I. A. Al-Shehbaz, A. Franzke, K. Mummenhoff, A. Stamatakis, M.A. Koch, BrassiBase: introduction to a novel knowledge database on Brassicaceae evolution. Plant Cell Physiol. 55, e3–e3 (2014). doi:10.1093/pcp/pct158

24. The Plant List Version 1.1. Published on the Internet (accessed 1st January 2021). http://www.theplantlist.org/1.1/browse/A/Brassicaceae/ (2013).

25. P. Legendre, L. Legendre, Numerical Ecology. Elsevier, Oxford (2012). ISBN:9780080523170

26. T. Guillerme, N. Cooper, S. L. Brusatte, K. E. Davis, A. L. Jackson, S. Gerber, A. Goswami, K. Healy, M. J. Hopkins, M. E. Jones, G. T. Lloyd, Disparities in the analysis of morphological disparity. Biol. Lett. 16, 20200199 (2020). doi:10.1098/rsbl.2020.0199

27. M. Méndez, J. M. Gómez, Phenotypic gender in *Hormathophylla spinosa* (Brassicaceae), a perfect hermaphrodite with tetradynamous flowers, is variable. Plant Syst. Evol. 262, 225–23 (2006). doi:10.1007/s00606-006-0462-5

28. V. L. Soza, V. Le Huynh, V. S. Di Stilio, Pattern and process in the evolution of the sole dioecious member of Brassicaceae. EvoDevo 5, 42 (2014). doi:10.1186/2041-9139-5-42

29. E. Narbona, H. Wang, P. L. Ortiz, M. Arista, E. Imbert, Flower colour polymorphism in the Mediterranean Basin: occurrence, maintenance and implications for speciation. Plant Biol. 20, 8–20 (2018). doi:10.1111/plb.12575.

30. C. A. Dick, J. Buenrostro, T. Butler, M. L, Carlson, D. J. Kliebenstein, J. B. Whittall, Arctic mustard flower color polymorphism controlled by petal-specific downregulation at the threshold of the anthocyanin biosynthetic pathway. PLoS One 6, e18230 (2011). doi:10.1371/journal.pone.0018230

31. B. Zhang, C. Liu, Y. Wang, X. Yao, F. Wang, J. Wu, G. J. King, K. Liu, Disruption of a carotenoid cleavage dioxygenase 4 gene converts flower colour from white to yellow in *Brassica* species. New Phytol. 206, 1513–1526 (2015). doi:10.1111/nph.13335

32. S. Faisal, Y. Guo, S. Zang, B. Cao, G. Qu, S. Hu, Morphological and genetic analysis of a cleistogamous mutant in rapeseed (*Brassica napus* L.). Genet. Resour. Crop Evol. 65, 397–403 (2018). doi:10.1007/s10722-017-0598-x

33. G. Theissen, Homeosis of the angiosperm flower: studies on three candidate cases of saltational evolution. Palaeodiversity 3, 131–139 (2010).

34. M. V. Byzova, J. Franken, M. G. Aarts, J. de Almeida-Engler, G. Engler, C. Mariani, M. M. V. L. Campagne, G. C. Angenent, *Arabidopsis* STERILE APETALA, a multifunctional gene regulating inflorescence, flower, and ovule development. Genes Dev. 13, 1002–1014 (1999). doi:10.1101/gad.13.8.1002.

35. T. Van der Niet, S. D. Johnson, Phylogenetic evidence for pollinator-driven diversification of angiosperms. Trends Ecol. Evol. 27, 353–361 (2012). doi:10.1016/j.tree.2012.02.002

36. A. Franzke, M. A. Lysak, I. A. Al-Shehbaz, M. A. Koch, K. Mummenhoff, Cabbage family affairs: the evolutionary history of Brassicaceae. Trends Plant Sci. 16, 108–116 (2011). doi:10.1016/j.tplants.2010.11.005

37. J. M. Gómez, F. Perfectti, J. Lorite, The role of pollinators in floral diversification in a clade of generalist flowers. Evolution 69, 863–878 (2015). doi:10.1111/evo.12632

38. S. Castiglione, C. Serio, D. Tamagnini, M. Melchionna, A. Mondanaro, M. Di Febbraro, A. Profico, P. Piras, F. Barattolo, P. Raia, A new, fast method to search for morphological convergence with shape data. PloS One 14: e0226949 (2019). doi:10.1371/journal.pone.0226949

39. C. T. Stayton, The definition, recognition, and interpretation of convergent evolution, and two new measures for quantifying and assessing the significance of convergence. Evolution 69, 2140–2153 (2015). doi:10.1111/evo.12729

40. K. Arbuckle, C. M. Bennett, M. P. Speed, A simple measure of the strength of convergent evolution. Methods Ecol. Evol. 5, 685–693 (2014). doi:10.1111/2041-210X.12195

41. K. Faegri, L. Van Der Pijl, Principles of Pollination Ecology. Elsevier, Oxford (1980). ISBN:9780080164212

42. A. S. Dellinger, Pollination syndromes in the 21st century: where do we stand and where may we go?. New Phytol. 228, 1193–1213 (2020). doi:10.1111/nph.16793

43. R. D. Phillips, R. Peakall, T. van der Niet, S. D. Johnson, Niche perspectives on plant– pollinator interactions. Trends Plant Sci. 25, 779–793 (2020). doi:10.1016/j.tplants.2020.03.009.

44. C. A. Wessinger, L. C. Hileman, Parallelism in flower evolution and development. Annu. Rev. Ecol. Evol. Syst. 51, 387–408 (2020). doi:10.1146/annurev-ecolsys-011720-124511

45. J. D. Thomson, P. Wilson, Explaining evolutionary shifts between bee and hummingbird pollination: convergence, divergence, and directionality. Int. J. Plant Sci. 169, 23–38 (2008). doi:10.1086/523361

46. J. M. Gómez, F. Perfectti, M. Abdelaziz, J. Lorite, A. J. Muñoz Pajares, J. Valverde, Evolution of pollination niches in a generalist plant clade. New Phytol. 205, 440–453 (2015). doi:10.1111/nph.13016

47. E. C. Snell-Rood, M. E. Kobiela, K. L. Sikkink, A. M. Shepherd, Mechanisms of plastic rescue in novel environments. Annu. Rev. Ecol. Evol. Syst. 49, 331–354 (2018). doi:10.1146/annurev-ecolsys-110617-062622

48. R. J. Fox, J. M. Donelson, C. Schunter, T. Ravasi, J. D. Gaitán-Espitia, Beyond buying time: the role of plasticity in phenotypic adaptation to rapid environmental change. Phil. Trans. R. Soc. B 374, 20180174 (2019). doi:10.1098/rstb.2018.0174

49. N. A. Levis, D. W. Pfennig, Plasticity-led evolution: evaluating the key prediction of frequency-dependent adaptation. Proc. Biol. Sci. 286, 20182754 (2019). doi:10.1098/rspb.2018.2754

50. M. R. Warner, L. Qiu, M. J. Holmes, A. S. Mikheyev, T. A. Linksvayer, Convergent eusocial evolution is based on a shared reproductive groundplan plus lineage-specific plastic genes. Nature Comm. 10, 1–11 (2019). doi:10.1038/s41467-019-10546-w

51. R. F. Schneider, A. Meyer, How plasticity, genetic assimilation and cryptic genetic variation may contribute to adaptive radiations. Mol. Ecol. 26, 330–350 (2017). doi:10.1111/mec.13880

52. T. Guillerme, N. Cooper, Time for a rethink: time sub sampling methods in disparity through time analyses. Palaeontology 61, 481–493 (2018). doi:10.1111/pala.12364

53. P. Legendre, D. Borcard, P. R. Peres-Neto, Analyzing beta diversity: Partitioning the spatial variation of community composition data. Ecol. Monogr. 75, 435–450 (2005). doi:10.1890/05-0549

54. P. Mair, I. Borg, Rusch T. Goodness-of-fit assessment in multidimensional scaling and unfolding. Multivar. Behav. Res. 51, 772–789 (2016). doi:10.1080/00273171.2016.1235966.

55. J. B. Kruskal, Multidimensional scaling by optimizing goodness-of-fit to a nonmetric hypothesis. Psychometrika 29, 1–28 (1964).

56. J. B. Kruskal, Nonmetric multidimensional scaling: a numerical method. Psychometrika 29, 115–129 (1964).

57. G. De’ath, Extended dissimilarity: a method of robust estimation of ecological distances from high beta diversity data. Plant Ecol. 144, 191–199 (1999). doi:10.1023/A:1009763730207

58. J. Oksanen, F. G. Blanchet, R. Kindt, P. Legendre, P. R. Minchin, R. B. O’hara, G. L. Simpson, P. Solymos, M. H. H. Stevens, H. Wagner, 2013 Package ‘vegan’. Community ecology package, version 2, 1–295 https://cran.r-project.org, https://github.com/vegandevs/vegan (2019)

59. S. C. Goslee, Urban, D. L The ecodist package for dissimilarity-based analysis of ecological data. J. Stat. Softw. 22, 1–19 (2007). doi:10.18637/jss.v022.i07

60. J. Oksanen, Vegan: an introduction to ordination. *URL* http://cran.r-project.org/web/packages/vegan/vignettes/introvegan.pdf (2020).

61. T. Guillerme, M. N. Puttick, A. E. Marcy, V. Weisbecker, Shifting spaces: Which disparity or dissimilarity measurement best summarize occupancy in multidimensional spaces?. Ecol. Evol. 10, 7261–7275 (2020). doi:10.1002/ece3.6452

62. T. Guillerme, dispRity: a modular R package for measuring disparity. Methods Ecol. Evol. 9, 1755–1763 (2018). doi:10.1111/2041-210X.13022

63. M. A. Koch, M. Kiefer, D.A. German, I.A. Al-Shehbaz, A. Franzke, K. Mummenhoff, R. Schmickl, BrassiBase: Tools and biological resources to study characters and traits in the Brassicaceae — version 1.1. Taxon 61, 1001–1009 (2012). doi:10.1002/tax.615007

64. D. J., Bennett, M. D., Sutton, S. T. Turvey, Treeman: an R package for efficient and intuitive manipulation of phylogenetic trees. BMC Res. Notes 10, 30 (2017). doi:10.1186/s13104-016-2340-8

65. M. A. Ragan, Phylogenetic inference based on matrix representation of trees. Mol. Phylogenetics Evol. 1, 53–58 (1992). doi:10.1016/1055-7903(92)90035-F

66. S. Bayat, M. E. Schranz, E. H. Roalson, J. C. Hall, Lessons from Cleomaceae, the sister of crucifers. Trends Plant Sci. 23, 808–821 (2018). doi:10.1016/j.tplants.2018.06.010.

67. C. Boettiger, D. Temple Lang, Treebase: an R package for discovery, access and manipulation of online phylogenies. Methods Ecol. Evol. 3, 1060–1066 (2012). doi:10.1111/j.2041-210X.2012.00247.x

68. E., Paradis, K. Schliep, ape 5.0: an environment for modern phylogenetics and evolutionary analyses in R. Bioinformatics 35, 526–528 (2019). doi:10.1093/bioinformatics/bty633

69. K. Schliep, phangorn: phylogenetic analysis in R. Bioinformatics 27, 592–593 (2011). doi:10.1093/bioinformatics/btq706

70. L. J. Revell, phytools: an R package for phylogenetic comparative biology (and other things). Methods Ecol. Evol. 3, 217–223 (2012). doi:10.1111/j.2041-210X.2011.00169.x

71. O. J. Hardy, S. Pavoine, Assessing phylogenetic signal with measurement error: a comparison of Mantel tests, Blomberg et al.’s *K*, and phylogenetic distograms. Evolution 66, 2614–2621 (2012). doi:10.1111/j.1558-5646.2012.01623.x

72. J. Clavel, G. Escarguel, G. Merceron, mvMORPH: an R package for fitting multivariate evolutionary models to morphometric data. Methods Ecol. Evol. 6, 1311–1319 (2015). doi:10.1111/2041-210X.12420

73. F. Perfectti, J. M. Gómez, A. González-Megías, M. Abdelaziz, J. Lorite, Molecular phylogeny and evolutionary history of *Moricandia* DC (Brassicaceae). PeerJ 5, e3964 (2017). doi:10.7717/peerj.3964

74. S. A. Smith, J. W. Brown, Constructing a broadly inclusive seed plant phylogeny. Am. J. Bot. 105, 302–314 (2018). doi:10.1002/ajb2.1019

75. M. L. Gaynor, J. Ng, R. G. Laport, Phylogenetic structure of plant communities: are polyploids distantly related to co-occurring diploids?. Frontiers Ecol. Evol. 6, 52 (2018). doi:10.3389/fevo.2018.00052

76. X.-C. Huang, D. A. German, M. A. Koch, Temporal patterns of diversification in Brassicaceae demonstrate decoupling of rate shifts and mesopolyploidization events. Ann. Bot. 125, 29–47 (2019). doi:10.1093/aob/mcz123

77. J. M. Gómez, M. Verdú, F. Perfectti, Ecological interactions are evolutionarily conserved across the entire tree of life. Nature 465, 918–921 (2010). doi:10.1038/nature09113

78. C. B. Fenster, W. S. Armbruster, P. Wilson, M. R. Dudash, J. D. Thomson, Pollination syndromes and floral specialization. Annu. Rev. Ecol. Evol. Syst. 35, 375–403 (2004). doi:10.1146/annurev.ecolsys.34.011802.132347

79. J. M. Gómez, R. Torices, J. Lorite, C. P. Klingenberg, F. Perfectti, The role of pollinators in the evolution of corolla shape variation, disparity and integration in a highly diversified plant family with a conserved floral bauplan. Ann. Bot. 117, 889–904 (2016). doi:10.1093/aob/mcv194

80. C. F. Dormann, R. Strauss, A method for detecting modules in quantitative bipartite networks. Methods Ecol. Evol. 5, 90–98 (2014). doi:10.1111/2041-210X.12139

81. R. Guimerà, L. A. N. Amaral, Functional cartography of complex metabolic networks. Nature 433, 895–900 (2005). doi:10.1038/nature03288

82. M. E. J. Newman, Analysis of weighted networks. Phys. Rev. E70, 056131 (2004). doi:10.1103/PhysRevE.70.056131

83. C. F. Dormann, B. Gruber, J. Fründ, Introducing the bipartite package: Analysing ecological networks. R News 8, 8–11 (2008).

84. P. Feinsinger, E. E. Spears, R. W. Poole, A simple measure of niche breadth. Ecology 62, 27–32 (1981). doi:10.2307/1936664

85. J. Zhang, M. J. Zhang, Package ‘spaa’. R package version 1. https://CRAN.R-project.org/package=spaa (2013).

86. J. P. Huelsenbeck, R., Nielsen, J. P. Bollback, Stochastic mapping of morphological characters. Syst. Biol. 52, 131–158 (2003). doi:10.1080/10635150390192780

87. J. P. Bollback, SIMMAP: stochastic character mapping of discrete traits on phylogenies. BMC Bioinformatics 7, 88 (2006). doi:10.1186/1471-2105-7-88

88. L. J. Harmon, J. T. Weir, C. D. Brock, R. E. Glor, W. Challenger, GEIGER: investigating evolutionary radiations. Bioinformatics 24, 129–131 (2008). doi:10.1093/bioinformatics/btm538

89. C. T. Stayton, Convevol: quantifies and assesses the significance of convergent evolution. R package version 1.0. See http://cran.r-project.org/web/packages/convevol/index.html (2018).

90. P. Raia, S. Castiglione, C. Serio, A. Mondanaro, M. Melchionna, M. Di Febbraro, RRphylo: Phylogenetic ridge regression methods for comparative studies. Methods Ecol. Evol. 9, 974–983 (2019). doi:10.1111/2041-210X.12954

91. K., Arbuckle, A. Minter, Windex: Analyzing convergent evolution using the Wheatsheaf index in R. *Evol*. Bioinformatics 11, EBO-S20968 (2015). doi:10.4137/EBO.S20968

